# The Proteomic Landscape of Genome-Wide Genetic Perturbations

**DOI:** 10.1101/2022.05.17.492318

**Authors:** Christoph B. Messner, Vadim Demichev, Julia Muenzner, Simran Aulakh, Annika Röhl, Lucía Herrera-Domínguez, Anna-Sophia Egger, Stephan Kamrad, Oliver Lemke, Enrica Calvani, Michael Mülleder, Kathryn S. Lilley, Georg Kustatscher, Markus Ralser

## Abstract

Functional genomic strategies help to address the genotype phenotype problem by annotating gene function and regulatory networks. Here, we demonstrate that combining functional genomics with proteomics uncovers general principles of protein expression, and provides new avenues to annotate protein function. We recorded precise proteomes for all non-essential gene knock-outs in *Saccharomyces cerevisiae.* We find that protein abundance is driven by a complex interplay of i) general biological properties, including translation rate, turnover, and copy number variations, and ii) their genetic, metabolic and physical interactions, including membership in protein complexes. We further show that combining genetic perturbation with proteomics provides complementary dimensions of functional annotation: proteomic profiling, reverse proteomic profiling, profile similarity and protein covariation analysis. Thus, our study generates a resource in which nine million protein quantities are linked to 79% of the yeast coding genome, and shows that functional proteomics reveals principles that govern protein expression.

**Highlights:**

- Nine million protein quantities recorded in ~4,600 non-essential gene deletions in *S. cerevisiae* reveal principles of how the proteome responds to genetic perturbation
- Genome-scale protein expression is determined by both functional relationships between proteins, as well as common biological responses
- Broad protein expression profiles in slow-growing strains can be explained by chromosomal aneuploidies
- Protein half-life and ribosome occupancy are predictable from protein abundance changes across knock-outs
- Functional proteomics annotates missing gene function in four complementary dimensions

## Introduction

Understanding how changes in the genotype cause a given phenotype is a key question of molecular biology and a challenge that is of shared importance for precision medicine, biotechnology, and synthetic biology. Predicting the phenotype of a mutant requires intricate knowledge of how protein networks respond to perturbation as well as a system-wide understanding of protein function. However, even in the best studied species, such as humans and budding yeast, a substantial fraction of the proteome remains without detailed functional annotation, while at the same time research efforts continue to focus on a small number of genes already known in the pre-genomic era (Kustatscher et al., 2022).

The biological consequences of a genetic perturbation can be complex. For instance, the loss of a protein may deprive a cell of one molecular function such as an enzymatic activity, but equally could have repercussions for a wide range of biological processes. If the protein is part of a complex, it can lead to its disintegration and the degradation of non-assembled subunits by non-exponential decay (McShane et al., 2016). The lack of an individual protein could disrupt metabolic pathways, reconfigure other pathways, or indirectly affect cell state (Deutscher et al., 2006; Liu et al., 2019; Stefely et al., 2016). Therefore, even the phenotype of single gene loss disease can be complex, a problem well known from monogenic disease (Weatherall, 2001). For example, triosephosphate isomerase deficiency is a rare human disease mapped in the 1960s, resulting from loss of function of a single glycolytic enzyme (*TPI1*) with a well understood function in a well understood pathway, glycolysis. Yet the aetiology of the syndrome is only partially understood to date, and the phenotypic spectrum associated with the enzyme deficiency ranges from hemolytic anaemia, muscle degeneration, immune defects, and up to mental retardation (Orosz et al., 2006; Segal et al., 2019). Discerning the principles of proteome remodelling following genetic perturbation could improve our understanding of protein interaction networks, the regulation of protein complexes, the evolution of protein expression, and eventually, how a phenotype emerges from a change in the genotype. In addition, mutations that lead to a loss of protein expression (or function) are common causes of drug resistance in microbes, fungal pathogens, and of malignant diseases. For example, genomic abnormalities are widespread in cancer cells, making it difficult to identify which mutations are key to driving cancer growth. Moreover, the relationship between oncogenic mutations and the therapeutically exploitable vulnerabilities they cause is poorly understood (Boehm et al., 2021). Hence, a genome-wide understanding of protein expression dynamics could facilitate the manipulation of metabolic and protein–protein interaction networks, a promising therapeutic opportunity (Hahn et al., 2021).

With the availability of genome-editing tools such as CRISPR-Cas9, that are more efficient than homologous recombination, systematic mutant libraries are increasingly becoming available for mapping both protein function as well as the consequences of genetic perturbation. If genome editing is applied at the genome scale, *functional genomic* experiments are facilitated. Using systematic knock-out libraries for functional genomic experiments was spearheaded by the yeast knock-out strain collection (Giaever and Nislow, 2014; Winzeler et al., 1999), which has opened fundamental avenues to understand genetic interactions, chemical–genetic interactions, drug resistance, as well as their impact on genome, transcriptome, metabolome, and phenome (Costanzo et al., 2010; Giaever et al., 2002; Hillenmeyer et al., 2008; Marguerat et al., 2012; Mülleder et al., 2016). Moreover, the combination of the library with transcriptomics and metabolomics provided new functional information for individual gene knock-outs, and enabled the characterisation of unknown genes using guilt-by-association approaches (Kemmeren et al., 2014; Mülleder et al., 2016).

Although proteins are the primary output of gene expression, the impact of systematic genetic perturbations on the proteome remains poorly understood. Indeed, until recently, it was challenging to apply proteome technologies at a very large scale. Proteomes were instead measured for specific subsets of the gene knock-out collection—for instance, focussing on mitochondrial function (Stefely et al., 2016), kinase activity (Zelezniak et al., 2018), or central carbon metabolism (Matsuda et al., 2017). These studies demonstrated a substantial added value for applying proteomics systematically to understanding both gene function and gene networks.

Recently, we and others have demonstrated that through the application of robust chromatographic regimes, new sample preparation strategies that reduce batch effects, and new acquisition schemes and software (Bache et al., 2018; Bekker-Jensen et al., 2020; Bian et al., 2020; Bruderer et al., 2019; Demichev et al., 2020; Geyer et al., 2016; Messner et al., 2021; Muenzner et al., 2022), proteomics can be scaled to thousands of samples, without compromising measurement precision. Herein, we apply high-throughput proteomics in order to understand the proteomic landscape of genome-wide genetic perturbations by measuring precise quantitative proteomes for all non-essential gene deletions in *Saccharomyces cerevisiae*. We thus create, to our knowledge, one of the largest and most systematic proteomic datasets for any species, resulting in a phenotypic map of almost nine million precise protein quantities which provide functional information for 80% of yeast genes, and that reveal general principles that underlie protein expression.

## Results

### Precise quantitative proteomes for gene knock-outs at a genome-wide scale

We systematically measured the yeast proteome upon the systematic knock-out of (non-essential) genes in the haploid yeast *Saccharomyces cerevisiae* strain BY4741, a derivative of the S228c background (Giaever and Nislow, 2014; Winzeler et al., 1999). To conduct our studies under conditions where a majority of metabolic pathways are active, we used a modified version of the knock-out collection with restored prototrophy (Mülleder et al., 2012) and grew cells in a defined synthetic minimal (SM) media without amino-acid and nucleobase supplementation (Mülleder et al., 2012, 2016). Strains viable under those conditions were distributed across 57 96-well plates (Figure 1A). To ensure measurement quality and reproducibility, we adapted the library for a large-scale omic analysis, and added 375 wild-type strain controls (WT; complemented *his3Δ* deletion strain). In addition, we measured 389 quality control (QC) samples (pooled yeast digest, 7 per plate) alongside the data acquisition, bringing it to a total of 764 proteomes measured as controls (Figure 1A).

**Figure 1.**
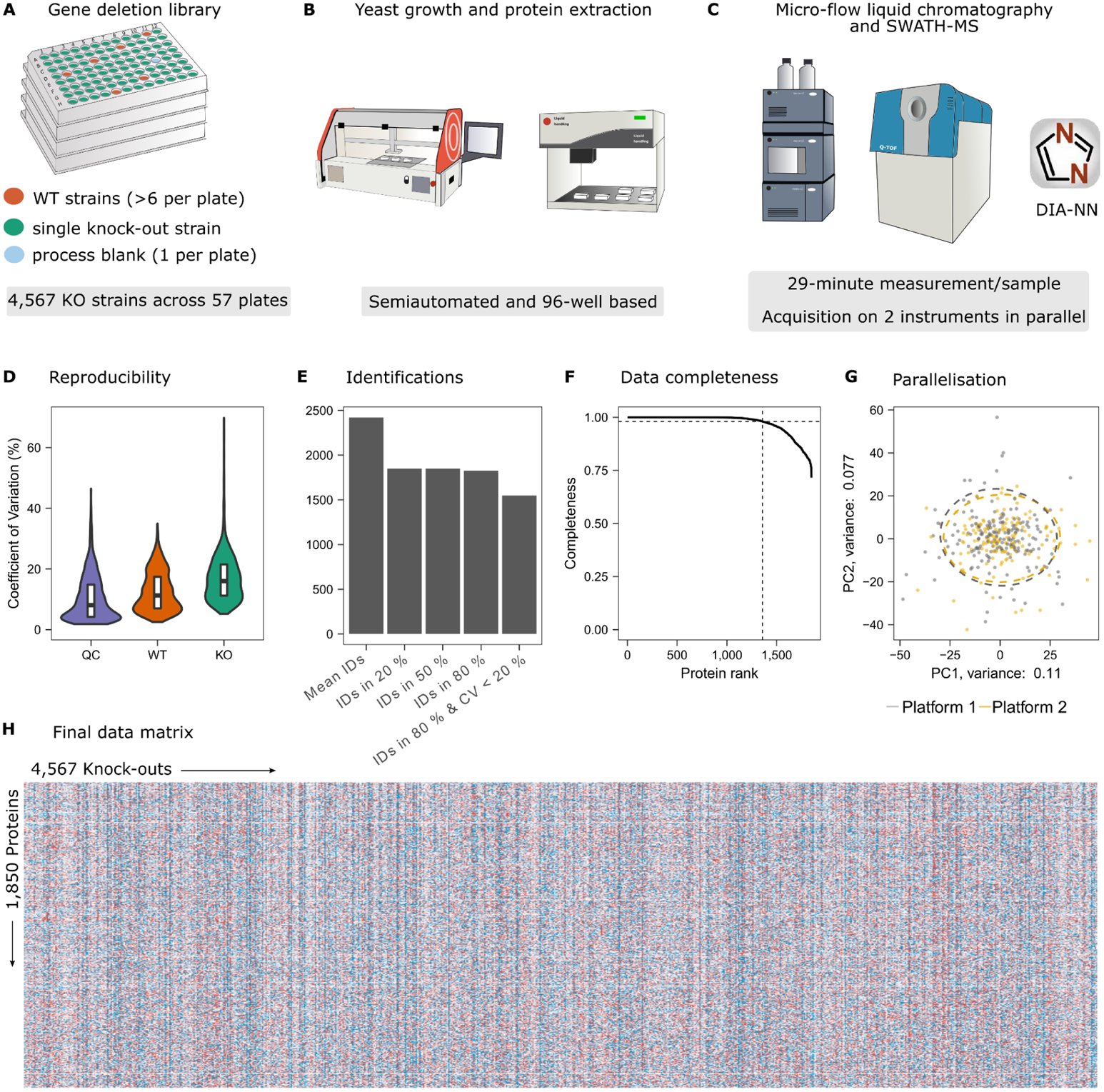
Precise quantitative proteomes for the genome-scale yeast gene-deletion collection in minimal medium. **(A) Library design.** A prototrophic version (Mülleder et al., 2012) of the haploid (MAT**a**) deletion collection (Winzeler et al., 1999) was distributed across 57 96-well plates, and complemented with at least 6 wild-type controls and one blank to monitor cell growth and analytical precision, on each plate. **(B) Streamlined sample preparation.** Liquid handling, as well as physical cell rupture, reduced hands-on time while allowing a throughput of 384 samples/day and reproducible and consistent sample processing across the large study (Methods). **(C) High-throughput proteomic data acquisition and analysis using an optimised microLC-SWATH-MS workflow.** The samples were measured on two identical LC-MS setups, consisting of microflow-rate liquid chromatography (nanoAcquity, Waters) and TripleTOF 6600 instruments (SCIEX). The peptides were separated with a 19-min non-linear gradient (~30 min total runtime; Table S2) using microflow (5 µl/min) and an adapted acquisition scheme using variable window SWATH (Muenzner et al., 2022). The raw data was processed by DIA-NN, which we provide as a free software to facilitate large scale proteomic DIA experiments using short gradient chromatography (Demichev et al., 2020). **(D) Precise proteome quantification and high data consistency in large-scale proteomic experiments.** The coefficients of variation (CV; in %) were calculated for each protein and are shown for pooled yeast digest samples (QC, n = 389), whole-process control samples (WT, n = 375), and KO samples (KO, n = 4,703). Median CV values are 8.1%, 11.3% capturing the technical variabilities, and 16.2%, capturing technical variability plus the biological signal, respectively. CVs were calculated on the filtered dataset (and are shown from 0 to 70 % (see Methods; see Figure S1C for all data points). **(E) Protein identifications**. Protein identification numbers are shown as mean per sample, identified in 20% of the samples, identified in 50% of the samples, identified in 80% of the samples as well as identified in 80% of the WT samples with CV < 20% (as described in Methods). **(F) Consistent identification of proteins.** Completeness was calculated for each protein as the number of samples in which the respective protein was identified divided by the total number of samples. The proteins were ranked by their completeness value (decreasing along x-axis). **(G) Robustness enables parallelisation across instruments.** A principal component analysis of all WT control samples (n = 375) shows no batch separation despite the use of the two different LC-MS platforms. **(H) Data matrix.** The acquired quantitative data are shown as a heatmap with 1,850 unique proteins measured across the 4,567 knock-outs, resulting in a total of 8,848,950 protein quantities.

To facilitate the generation of thousands of (yeast) proteome profiles, we have recently generated a high-throughput proteomics pipeline that uses a semi-automated sample-preparation workflow, microflow-SWATH-MS, and DIA-NN analysis (Demichev et al., 2020; Gillet et al., 2012; Muenzner et al., 2022). Herein, we further streamlined the procedures for reaching the scale of the yeast KO collection without compromising data quality (Figure 1B and 1C). The adaptations included the optimisation of yeast growth to ensure that typical deletion strains are in a comparable growth phase, and technical aspects of sample preparation, data acquisition and processing in order to minimise batch effects. For instance, we used pre-prepared, frozen stock-solution plates throughout the experiment to avoid reagent-related batch effects (see Methods). We then decided on a robust microflow (5 µl/min) liquid chromatography (LC) method with an active chromatographic gradient of 19 minutes and a total time from injection to injection of ~30 minutes (see Methods). The robustness of the developed workflow enabled the acquisition on two instruments in parallel without compromising data comparability (Figure 1G and Figure S1B). The 8 TB raw data were processed with DIA-NN 1.7.12 (Demichev et al., 2020) using a spectral library generated with gas-phase fractionation (Messner et al., 2021). The average number of quantified proteins per run was 2,421 (from 20,065 measured precursors), which is about half of the yeast proteome that is detectable when maximising proteomic depth in low-throughput experiments through extensive pre-fractionation and nano-flow rate chromatography (de Godoy et al., 2006). After stringent filtering, we quantified 1,850 unique proteins across the 4,567 measured knock-outs (Figure 1E), resulting in a total of 8,848,950 protein quantities (Figure 1H). The dataset has a low number of missing values and high data completeness with 1,358 proteins quantified in 98% of all samples (Figure 1F). The average of the obtained intensities correlates well with absolute protein copy numbers per cell obtained by stable-isotope-labelled internal standards (Lawless et al., 2016) (r = 0.75; Figure S1A), supporting the quantitative nature of the proteomic data. In the final dataset, the median protein coefficient of variation (CV) was 8.1% for pooled yeast digest samples (n = 389; instrument variability) and 11.3% for the WT replicates (n = 375; whole-workflow variability). The technical variability was much lower than the biological responses, indicated by higher CV values in the knock-outs compared to the control samples (16.2% for KOs) (Figure 1D and Figure S1C). Hence, our workflow generated a proteome for those non-essential gene knock-outs in *S. cerevisiae* which grow in minimal medium, at a high measurement precision, and with low numbers of missing values.

### The dominating cellular processes that drive differential protein expression upon genetic perturbation

We report that systematic gene deletion and proteomics are highly complementary with respect to which genes they cover (Figure 2A). For example, proteome profiling is biassed towards abundant proteins, whereas gene deletion against essential genes, that tend to be abundant and are readily detected by proteomics. Neither technique appears to be biassed by gene/protein length. Uncharacterised genes, as defined by YeastMine (Balakrishnan et al., 2012), are more likely to encode low abundant proteins and are thus covered via their deletion (Figure 2A). As a result of these technological complementaries, combining genetic perturbation and proteomics enables a more comprehensive coverage of the genome for functional analysis than either approach could on its own. In total, we address the function of ~5,300 yeast genes (Figure 2B), corresponding to 78% of the entire coding genome (Cherry et al., 2012). This is more than what has been detected with proteomics, even after extensive pre-fractionation (de Godoy et al., 2008; Webb et al., 2013), and also, a higher number than unique gene deletions that grew in minimal nutrient medium (Mülleder et al., 2016)

**Figure 2.**
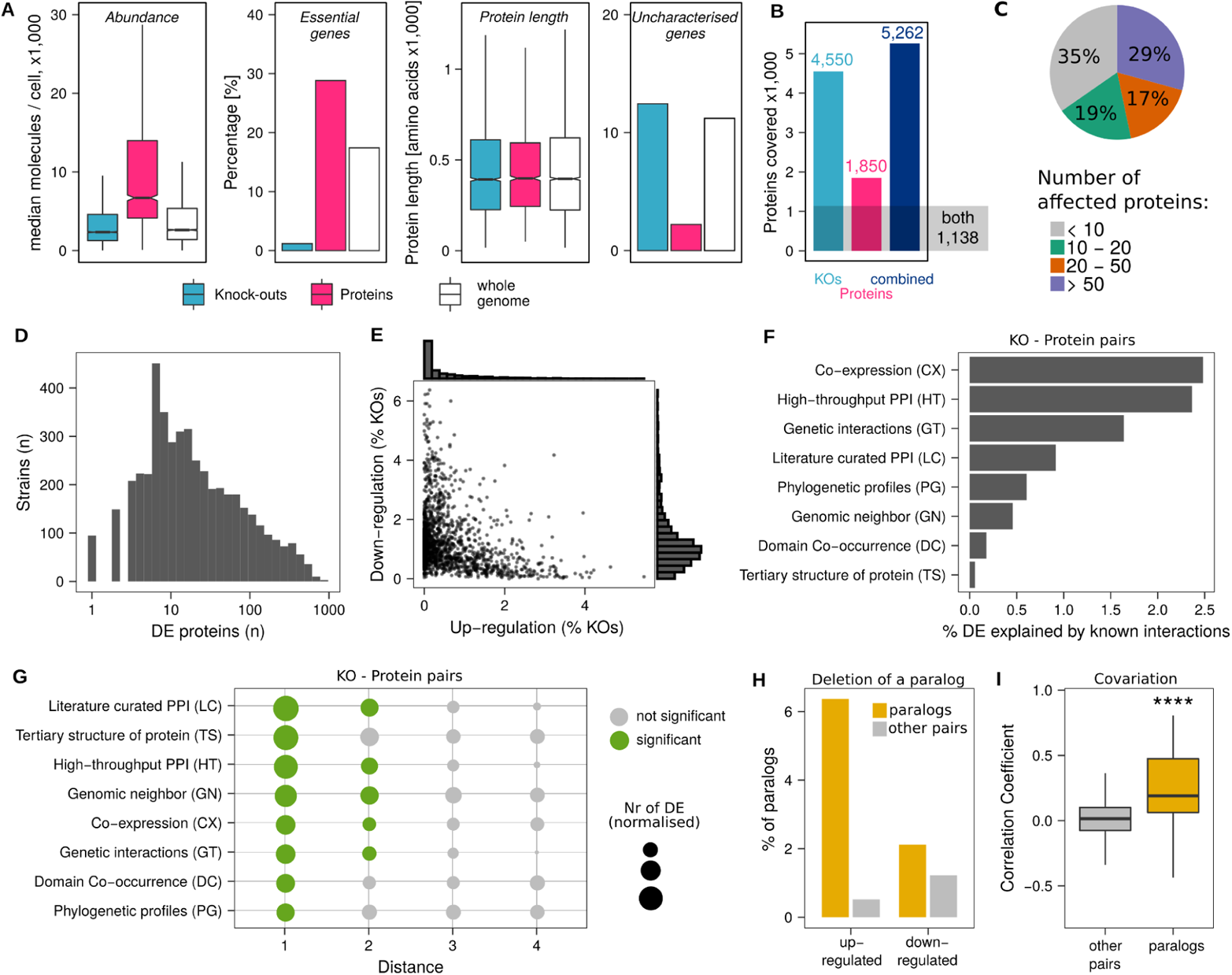
The proteomic response to systematic gene deletion reveals principles of protein expression. **(A)** Gene deletions and high-throughput proteomes capture complementary sets of proteins. Compared to the whole yeast genome, genes covered by mass spectrometry are biassed towards essential genes encoding for more-abundant proteins, whereas genes deleted in the KO library are more likely covering non-essential, more likely low abundant proteins that contain also more uncharacterised genes. **(B)** In combination, gene deletions and high-throughput proteomics cover 5,262 unique genes, which is 79% of the yeast genome annotation, and more than can be assessed with either technique in separation. 1,138 genes are covered by both knock-out and protein quantification. **(C) Responsiveness of the proteome to the genome-wide knock-outs.** Fraction of gene deletions (in %, total = 4,567) that cause a proteomic response of i) less than 10 proteins, ii) between 10 and 20, iii) between 20 and 50, or iv) more than 50 proteins (adjusted p-value < 0.01, BH for multiple testing correction (Benjamini and Hochberg, 1995)). **(D) Distribution of proteomic responses,** given as number of differentially expressed proteins (adjusted p-value < 0.01, BH for multiple testing correction). The x-axis is log_10_ transformed. **(E) Most proteins are either up-regulated or down-regulated, few proteins respond in both directions.** Up-regulations (x-axis) and down-regulations (y-axis) of each protein across the 4,567 KO strains are given as dots and as histograms. Numbers are given in % (differential regulation of a particular protein across the KO / total number of KO × 100). **(F) Physical, genetic, or functional interactions between genes explain 1/10th of differential protein expression**. Differentially expressed proteins (p-value < 0.01, BH for multiple testing) upon gene deletions were compared to physical, genetic or functional interactions, collected as part of the YeastNet ressource (v3, (Kim et al., 2014) Numbers are given in % (DE proteins explained by respective networks / total number of DE × 100). Some interactions are represented in more than one network, but the average overlap between two networks is less than 10% (Figure S2D). **(G) The effect of a deletion on a protein decreases with the distance to the knock-out gene in a physical, genetic or functional network.** The number of differential protein expressions (p-value < 0.01, BH for multiple testing) within the respective distances from the knock-out is indicated by dot size. Differentially expressed proteins of distance *i* were normalised to the total number of proteins of distance *i* within the respective network. Significance (hypergeometric test) is indicated by colour. (**H**) **Impact of deletions on paralogues.** Percentage of paralogues that are up- or down-regulated (adjusted p-value < 0.01, BH for multiple testing correction) after deletion of one of the paralogue partners (green). The number of up- and down-regulations across all KOs is shown as a grey bar for reference. **(I) Covariation of paralogs.** Correlation coefficients (Spearman) are shown for protein paralogues that arose from whole-genome duplication (Byrne and Wolfe, 2005) (n = 107 pairs) and for all other protein pairs (no paralogue, n = 1,710,215 pairs). The median Spearman correlation coefficients are 0.19 and 0.01 for paralogues and for all other pairs, respectively. Significance (Wilcoxon signed-rank test; **** for p-value ≤ 0.0001) is indicated with asterisks; the first and third quartiles, as well as the median (thick line), are shown with boxplots; whiskers extend to the most extreme data point that is no more than 1.5× the interquartile range from the box.

The number of proteins significantly affected by any gene deletion ranges up to 869 (p-value 0.01, BH for multiple testing (Benjamini and Hochberg, 1995)). 65% of the non-essential genes deleted cause the differential expression of > 10 proteins, 46% more than 20, and 29% more than 50 proteins (Figure 2C and 2D). Many profiles indicated a specific proteomic response to the gene deletion. For example, the knock-out strain *ARG81*, a transcription factor that represses arginine anabolism (Messenguy and Dubois, 2000), induces the differential expression of specifically 11 proteins, including Arg8, Arg3, Arg5, Arg56, and Arg1—the enzymes participating in the arginine biosynthetic pathway (Figure S2A). On the other hand, the knock-out of Rps27b, a protein of the small ribosomal subunit (40S), causes differential expression of 91 proteins. This broad profile of the *rps27bΔ* strain is the combination of proteins that are functionally related to Rps27b, but we also find a broad set of unrelated proteins differentially expressed. These are likely differentially expressed (DE) due to translation defects caused by the *rps27bΔ* deletion (Figure S2A). In general, gene knock-outs that directly or indirectly perturb translation or transcription by having Gene Ontology (GO) annotations such as *ribosomal small subunit progenesis, transcription from RNA polymerase I promotor or DNA-templated transcription, termination* have the strongest proteome responses, affecting the highest number of proteins (Figure S2G).

### The combination of gene knock-outs and proteomics reveals biological principles of gene expression

We found that upon genetic perturbation proteins are more often down-regulated than up-regulated. Each protein is on average up-regulated in 0.5% of all KOs (24 strains) while down-regulated in 1.2% of all KOs (55 strains). Furthermore, we were surprised to find that any individual protein changes predominantly in one direction (i.e. is either up-regulated or down-regulated) across the KO strains (Figure 2E). For example, Tsl1 or Tps2, both subunits of the trehalose-6-P synthase, are down-regulated in > 300 knock-outs while being up-regulated only in a few strains (Figure S2B). On the other hand, the tRNA synthetases Krs1, Hts1, or Frs1 are primarily up-regulated (Figure S2B).

Comparing the proteomic profiles to functional networks revealed that less than 10% of differential protein expression can be explained by the physical or functional interaction networks as assembled in YeastNet (Kim et al., 2014) (Figure 2F). For example, 2.4% of the differentially expressed proteins are connected with the knocked-out gene in a coexpression network or 2.3% in a high-throughput protein-protein interaction network (Figure 2F). After normalising to the number of interactions in the respective networks, we found that across different functional networks, proteins are significantly more likely differentially expressed if in direct neighbourhood to the deleted protein (Figure 2G). Only in some instances are secondary interactions also significantly increased, with the effect of the deletion decreasing with the distance to the gene knock-out (Figure 2G). For instance, direct interactions as well as secondary interactions in a literature-curated PPI network or in a genomic neighbour network have a significantly increased probability to be differentially expressed upon gene deletions.

We next investigated the interdependency of paralogues that arose by whole-genome duplication (ohnologues) (Byrne and Wolfe, 2005)). We found that in 8% of the cases where a paralogue was deleted, the other paralogue was up-regulated (Figure 2H, Table S5). The interdependence of paralogues was particularly evident for many ribosomal proteins such as Rpl16b, Rpl6a, or Rpl7a. Further, we detect a high level of co-expression for part of the ohnologues (21% with r > 0.5) (Figure 2I). For instance, *DCS1* / *DCS2* (r = 0.79) or *RNR2* / *RNR4* are co-regulated across the KO dataset (r = 0.8) (Figure S2E and S2F). Our data hence provide differentiated pictures about the role of paralogues that emerged from whole-genome duplication. For a fraction of these homologues, the proteome provides evidence for retained functional or regulatory relationships, because they are either co-regulated or differentially expressed when the other paralogues are missing. A majority of paralogues, however, are not co-expressed or do not respond to the genetic perturbation of the paralogue, indicating diversification of their function and/or their regulatory mechanisms.

### Mapping a complex relationship of growth rate, broad proteomic changes, and genome versatility

We detected less than 10% of differential protein expression in genes connected to the deleted gene via known genetic or physical interactions (Figure 2F). While this could be an underestimation given that such networks are still incomplete, this result highlights the role of general biological mechanisms that may guide differential protein expression. One known such factor is the growth rate, known to be correlated with changes in proteome and transcriptome (Airoldi et al., 2009; Fazio et al., 2008; Hughes et al., 2000a; Kemmeren et al., 2014; Kleijn et al., 2022; Slavov and Botstein, 2011; Wytock and Motter, 2019; Yu et al., 2021). Here, our large-scale dataset provides a differentiated picture about the role of growth-rate-dependent changes in differential protein expression. In agreement with previous studies, we find that slow-growing strains have a higher number of differentially expressed proteins and broader proteomic profiles (Figure 3A). Among those, strains with deletions of ribosome subunits have the slowest average growth rates (Figure S3A) and broad proteome responses with high numbers of differential expression (Figure S2G). Notably, however, our data showed that growth-rate-associated proteins explain only a fraction of differential protein expression in slow growers. For some of the proteins, we found moderate correlation with growth rate across the dataset (150 proteins have |r| > 0.4) (Figure 3C) and proteome expression changes allow growth rate predictions (Figure 3B). Indeed, a large fraction of differential expressions across the KOs are due to proteins that are not correlated with growth rate (Figure S3C). In the search for mechanisms that could explain broad responses of non-growth-correlated proteins in slow-growing strains, we realised that one key source of these profiles are aberrant chromosome numbers (aneuploidies). Aneuploidies are a common confounder to both growth and the proteome, as they cause broad expression changes by affecting all proteins encoded on an aneuploid chromosome (Torres et al., 2007). In lab strains, aneuploidies are transmitted to transcriptome and proteome with a minimum amount of gene-dosage buffering, and as a consequence are identifiable by proteomics similarly well as with genomics or transcriptomics, respectively (Muenzner et al., 2022; Pavelka et al., 2010; Torres et al., 2007). We sorted our protein-expression profiles according to the chromosomal localisation of the encoding genes, and analysed their expression patterns (Muenzner et al., 2022). We identified 110 aneuploid strains in our dataset (Figure 3D). For instance, the proteome of the deletion strain for the cell-cycle protein kinase gene *DBF2* reveals duplicated gene doses specifically for proteins encoded on chromosome VIII (Figure 3E). Segmental aneuploidies or short structural aneuploidies were detected in the proteome profiles of a further 18 strains, often in conjunction with whole-chromosome aneuploidies (Figure 3D). For instance, deletion for the spindle pole body component *KRE28*, which carries whole-chromosome aneuploidies on chromosomes II and VIII, as well as a segmental aneuploidy on chromosome VII (Figure 3F). Chromosomes IX, VIII, V, and I are changing their copy number most often, and we observed all chromosomes except for VI and VII to be aneuploid at least once (Figure S3F). Aneuploidies on these two chromosomes might be detrimental, and indeed isolated aneuploidies of chr VI were previously reported to be lethal due to genes such as alpha tubulin (*TUB2*) being located on that chromosome (Chan and Botstein, 1993).

**Figure 3.**
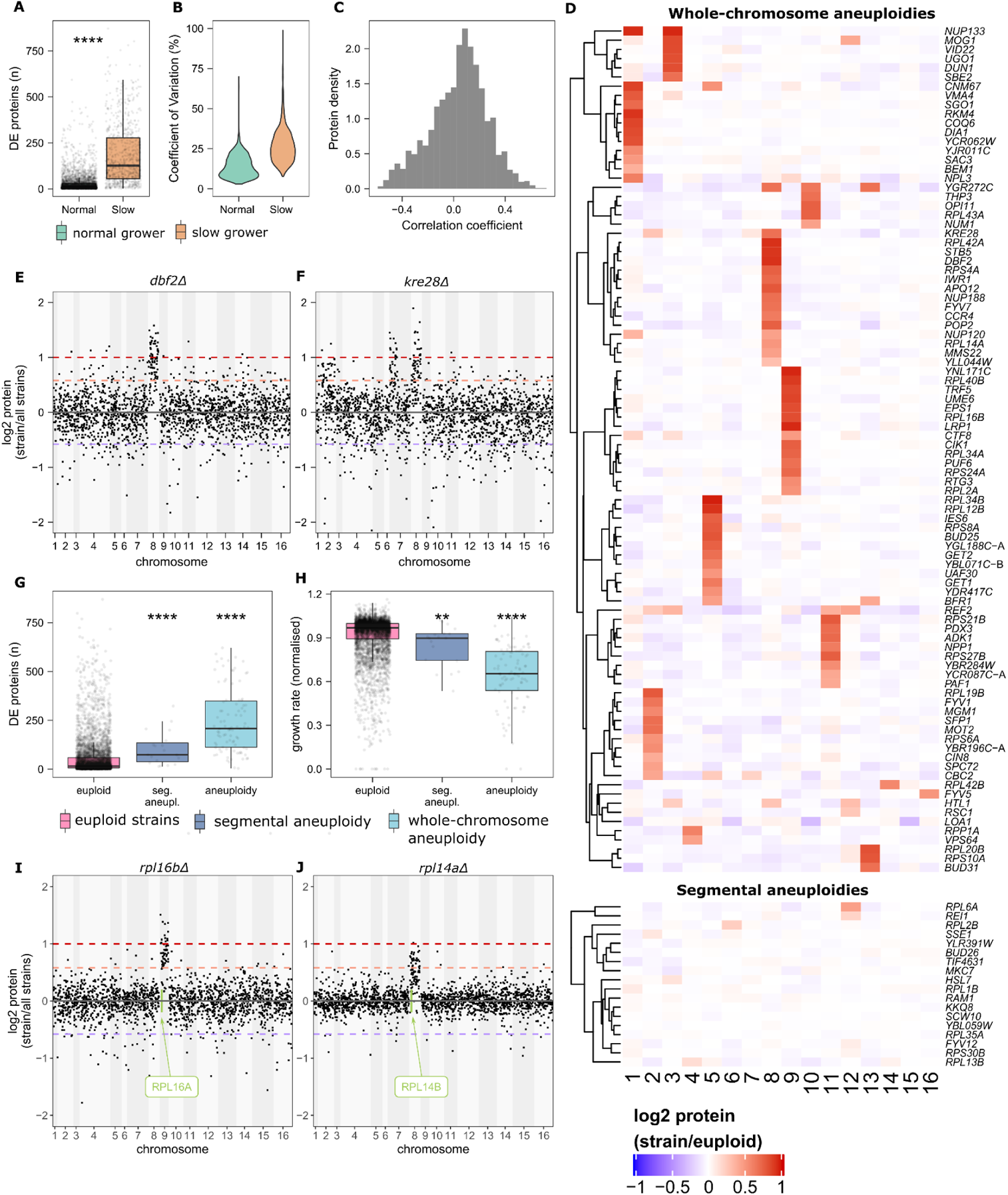
Slow-growing cells have broad proteomic changes of non-growth-related proteins, in part due to chromosomal copy-number alterations. **(A)** The numbers of differentially expressed proteins (adjusted p-value < 0.01, BH for multiple testing (Benjamini and Hochberg, 1995)) are compared between slow growers and normal growers. Slow growers were defined as normalised growth rate < 0.8 (n = 748) and normal growers ≥ 0.8 (n = 3,930). Significance (Wilcoxon signed-rank test; **** for p-value ≤ 0.0001) is shown with asterisks. **(B) Proteome variability among slow-growing strains.** The proteome dispersion within slow-growing strains is compared to the dispersion within normal-growing strains and is given as protein coefficients of variations (in %). Slow growers and normal growers were defined as normalised growth rates between 0.3 and 0.4; and normal-growing strains between 0.9 and 1.0. The CV values are shown for CV < 100%. **(C) Correlation of protein abundances with growth rate.** Correlation coefficients (Pearson correlation) are shown as histograms for all pairwise protein-abundance–growth correlations. **(D) Aneuploidy signatures across KO strains.** For each strain, log_2_ ratios between protein abundances and the median expression of the respective protein across all strains (presumed euploid) were calculated. Median log_2_ protein expression levels are shown for each chromosome. Strains are grouped into whole-chromosome aneuploidies and segmental aneuploidies and are labelled with the knocked-out gene name. **(E / F) Examples of aneuploid strains identified by proteomics.** Protein abundances relative to their median expression across all strains are shown for *dbf2Δ* and *kre28Δ,* respectively (Manhattan plot). Proteins were sorted by chromosomal location. The dashed lines indicate relative expressions of 2 (duplication) and 1.5. (**G**) **Aneuploid strains have high numbers of differentially expressed proteins.** The numbers of significantly changed proteins (adjusted p-value < 0.01, BH for multiple testing correction) are compared between euploid (n = 4,428, median = 16), segmental aneuploidy (n = 18, median = 74), and whole-chromosomal aneuploidy strains (n = 84, median = 208). The first and third quartiles, as well as the median (thick line), are shown with boxplots; whiskers extend to the most extreme data point that is no more than 1.5× the interquartile range from the box. **(H) Aneuploidy results in reduced growth rates.** The normalised growth rates are compared between euploid (n = 4,428, median = 0.97), segmental aneuploidy (n = 18, median = 0.90), and whole-chromosomal aneuploidy strains (n = 84, median = 0.65). The first and third quartiles, as well as the median (thick line), are shown with boxplots; whiskers extend to the most extreme data point that is no more than 1.5× the interquartile range from the box. **(I / J) Examples of knock-outs that cause aneuploidy for the paralogue-containing chromosome.** Protein abundances, sorted by their chromosomal location, are shown for *rpl16bΔ* and *rpl14aΔ,* respectively. Protein intensities were log_2_-transformed and normalised to their median expression levels across all strains. The dashed lines indicate relative expressions of 2, 1.5 and 0.67.

Our data confirm that aneuploidy is a cause of broad, non-growth-related proteomic responses in slow-growing strains. First, as in laboratory-engineered aneuploids (Pavelka et al., 2010; Torres et al., 2007), the aneuploids detected by our approach had a slow growth rate (Figure 3H). Second, these strains had broader, unspecific proteomic profiles (Figure 3G). For example, the aneuploid strain *kre28Δ* results in differential expression of 359 proteins, which is among the strongest proteome responses measured. Thus, in these strains, the slow growth rate and the broad proteome profile are not due to growth-rate-related protein changes, but explained by a common confounder: aberrant chromosome numbers.

We next asked whether there is a functional relationship between the deleted gene and the proteomic response induced by the aneuploidy. Overall, we found that aneuploid strains are enriched for gene deletions in ribosomal proteins as well as proteins involved in cell cycle and transcription. The specific enrichment of functional terms indicates that aneuploidies emerged as a response to the primary perturbation (Figure S3D). In agreement with early expression-profiling work (Hughes et al., 2000b) and more recent whole-genome sequencing data (Puddu et al., 2019), we found that knock-outs of ribosomal subunits, often encoded by two near-identical paralogues (Puddu et al., 2019), explain 17 out of 18 such compensatory chromosomal duplications. Further, our proteomic data show that in many cases the aneuploidy causes the up-regulation of a paralogue to the deleted gene (Figure S3E). For example, *rpl16bΔ* or *rpl14aΔ* cause aneuploidies of the chromosome IX and chromosome VIII, respectively, which contain their paralogues (Figure 3I and 3J). The expression levels of Rpl16a and Rpl14b are increased by fold-changes of 2.15 (adjusted p-value = 5.7 × 10^−46^) and 1.77 (adjusted p-value = 2.6 × 10^−6^), respectively. Thus, proteomic data suggest that chromosomal duplication can be a mechanism for ribosomal KOs for counteracting potential detrimental effects of gene knock-outs.

### Differential protein expression is dependent on translation rates and protein half-lives

We next addressed the role of protein dynamic factors, such as translation rate and protein turnover, in differential protein expression. We generated an elastic net regression model that aims to determine the translational profile as well as the protein turnover rate as a function of protein abundance changes in response to genetic perturbations. We used reference datasets for ribosome occupancy, determined by ribosomal profiling (McManus et al., 2014), as well as for protein turnover, obtained by metabolic labelling (Martin-Perez and Villén, 2017). We found that ~60% of the variation in ribosome occupancy is predicted from the proteome by the elastic net (Figure 4A), suggesting a strong link between translation rates and differential protein expression. Using elastic net feature weights, we assessed which KO strains were most informative for predicting ribosome occupancy (see Table S3 for a list of the ranked feature importance). The top weighted features were enriched for processes related to RNA levels or transcription (*mRNA processing, DNA-templated transcription, RNA splicing, transcription from RNA polymerase II promoter*) or protein degradation (*proteolysis involved in cellular protein catabolic process, protein modification by small protein conjugation)* (Figure 4B). This shows the interconnectivity of transcription, translation and degradation in the regulation of protein homeostasis.

**Figure 4.**
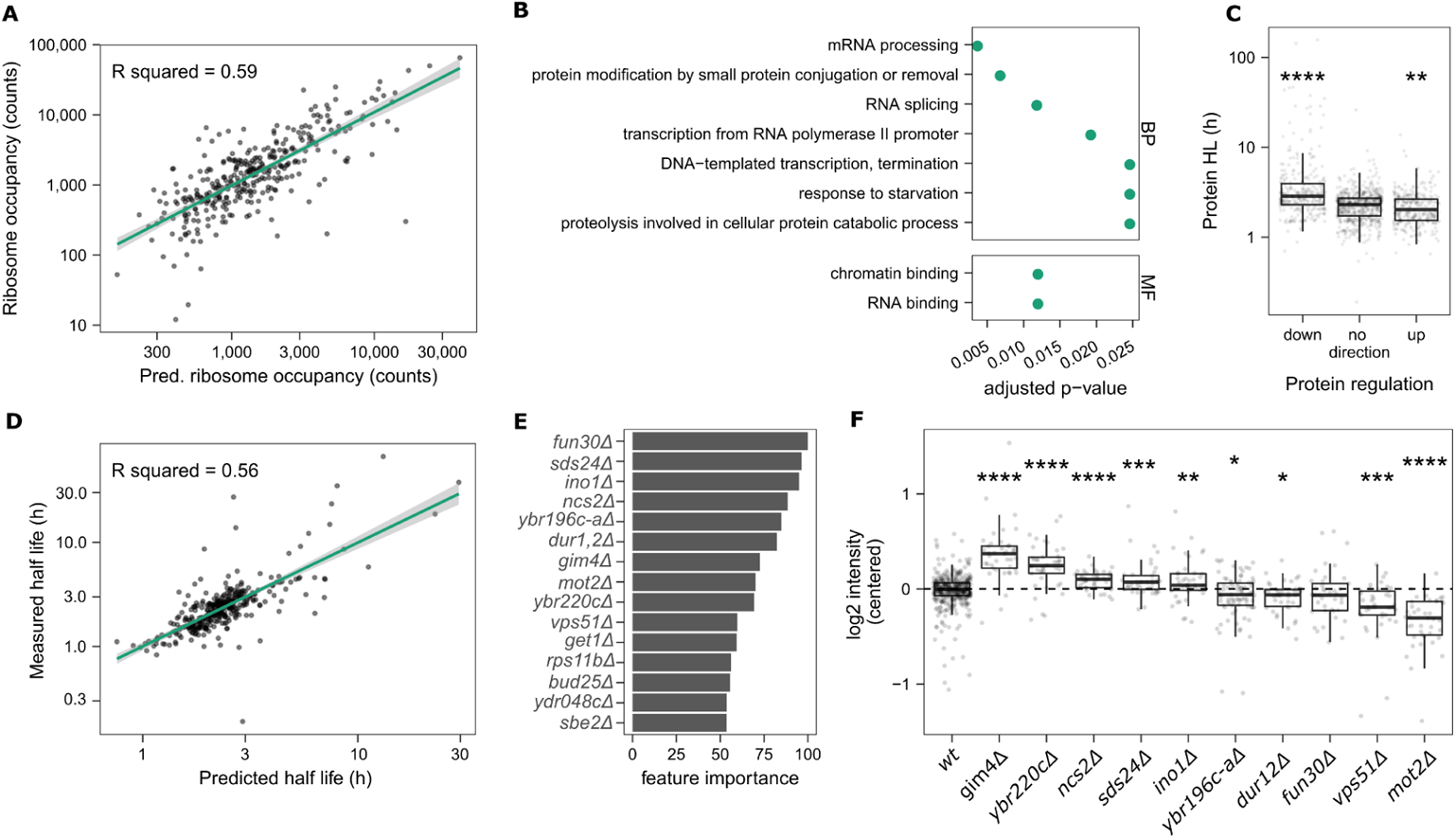
The interdependency of differential protein expression with translation rate and turnover. **(A) Machine learning predicts ribosomal occupancy from proteomics data.** An elastic net model with 10-fold cross-validation was trained on 80% of the data (n = 1,392) and plotted for the remaining 20% of the data (test set, n = 346). The proteome data was log_2_-transformed, centred, and scaled. Ribosomal occupancies were taken from a reference dataset (McManus et al., 2014) and log_10_-transformed. **(B)** Gene Ontology GO slim terms (Cherry et al., 2012)) enrichment among the top features selected by the model was conducted using hypergeometric testing. **(C) Proteins with long half-lives are more likely to be down-regulated**. Half-lives (in h) are shown as boxplots for proteins that are predominantly down-regulated, up-regulated, or change in both directions across the KO strains. Directionality was defined as ratios of up-regulation and down-regulation being > 75 % and < 25% quantile for down and up, respectively. Significance (two-sided Wilcoxon signed-rank test with “no direction” as a reference) is shown with asterisks (**** for p-value ≤ 0.0001; ** for p-value ≤ 0.01). The first and third quartiles, as well as the median (thick line), are shown with boxplots; whiskers extend to the most extreme data point that is no more than 1.5× the interquartile range from the box. **(D) Machine learning predicts protein half-life from proteomics data.** An elastic net model with 10-fold cross-validation was trained on 80% of the data (n = 1,398) and plotted for the remaining 20% of the data (n = 348). The proteome data was log_2_-transformed, centred, and scaled. Half-lives were taken from a reference dataset (Martin-Perez and Villén, 2017) and log_10_-transformed. **(E)** The 15 most important KO strains for the predictive model of half-lives. The KO strains are ranked by importance and scaled to have a maximum value of 100. **(F)** Abundance of ribosomal proteins (Rpl28, Rpl5, Rpl16b, Rpl16a, Rpl3, Rpl6a, Rpl6b, Rpl26b, Rpl26a, Rpl10, Rpl13b, Rpl32, Rpl8b, Rpl8a, Rpl25, Rpl37a, Rpl9a, Rpl9b, Rpl38, Rpl33a, Rpl33b, Rpl37b, Rpl36b, Rpl36a, Rpl22a, Rpl31a, Rpl30, Rpl24a, Rpl24b, Rpl39, Rpl7b, Rpl7a, Rpl4b, Rpl4a, Rpl29, Rpl14a, Rpl14b, Rpl17b, Rpl15a, Rpl17a, Rpl31b) in 10 KO strains that were selected as the most important feature for the prediction of protein half-life. Protein intensities are centred and log_2_-transformed. Significance for the comparison to the WT expression levels (two-sided t-test) is shown with asterisks (**** for p ≤ 0.0001; *** for p ≤ 0.001; ** for p ≤ 0.01; * for p ≤ 0.05; ns for p > 0.05). The first and third quartiles, as well as the median (thick line), are shown with boxplots; whiskers extend to the most extreme data point that is no more than 1.5× the interquartile range from the box.

Notably, our data also indicates a strong interdependence of protein turnover and the likelihood of a protein to be differentially expressed. We revealed that proteins with a slow turnover are more likely to be differentially expressed and tend to be down-regulated (Figure 4C). For example, the proteins Sds24, Hsp26, and Pgm2, which are among the most long-lived proteins in yeast (half-lives > 130 h), are primarily down-regulated (Figure S2C). Further, protein half-life was predictable by the elastic net from our proteomes (Figure 4D). Among the most determinant features (knock-outs) selected by the model we found a diverse set of genes such as *dur12Δ* (urea amidolyase), *sds24Δ* (protein involved in cell separation), and *fun30Δ* (involved in chromatin remodelling) (Figure 4E and Table S4), indicating that a variety of processes are directly or indirectly related to protein turnover. Their deletion results in a dependence of the expression levels on the protein half-lives (e.g. *dur12Δ* up-regulates long-lived proteins while *fun30Δ* down-regulates long-lived proteins) (Figure S4A). While neither growth rate nor cell size are the main driver of protein-half-life-dependent protein changes (Figure S4B), the translation machinery is significantly affected in the majority of those strains, indicated by up- or down-regulated ribosomal subunits (Figure 4F). Hence, our data establishes that protein abundance, translation rate, turnover are not independent properties, but rather interdependent and act together in determining the differential expression of a protein.

### The disruption of protein complexes can lead to the accelerated degradation of the surplus subunits, but also to their induction when feedback loops are involved

Our dataset allowed us to study the perturbation of all non-essential protein complex subunits in a single study. Protein complexes are stable assemblies of two or more physically interacting proteins that form functional modules with a defined stoichiometry. It is assumed that many complex subunits are produced in super-stoichiometric amounts and that excess subunits that are not incorporated into the complex (orphan proteins) are eventually degraded through non-exponential decay (NED) (Buccitelli and Selbach, 2020; Juszkiewicz and Hegde, 2018; McShane et al., 2016). We asked to which degree NED affects protein complexes disrupted on the genome-wide scale (Figure 5A). In 53% of the tested complexes, at least one of the knock-outs causes the significant differential expression of another complex subunit (adjusted p-value < 0.05, BH for multiple testing correction (Benjamini and Hochberg, 1995)) (Figure 5B and Figure S5). Out of those, 29% show down-regulation of the respective complex subunits (Figure 5B). For example, the knock-out of the gene *SEC28*, where the gene product has a stabilising function within the coatomer complex (Duden et al., 1998), induces down-regulation of its interacting subunits (Figure 5C). Other examples of down-regulation of affected subunits are the EGO complex (subunits Meh1 and Slm4 (Meldal and Orchard, 2018; Meldal et al., 2015, 2019)), or the alpha-1,6-mannosyltransferase complex (subunits Hoc1 and Mn11) (Figure S5-b).

**Figure 5.**
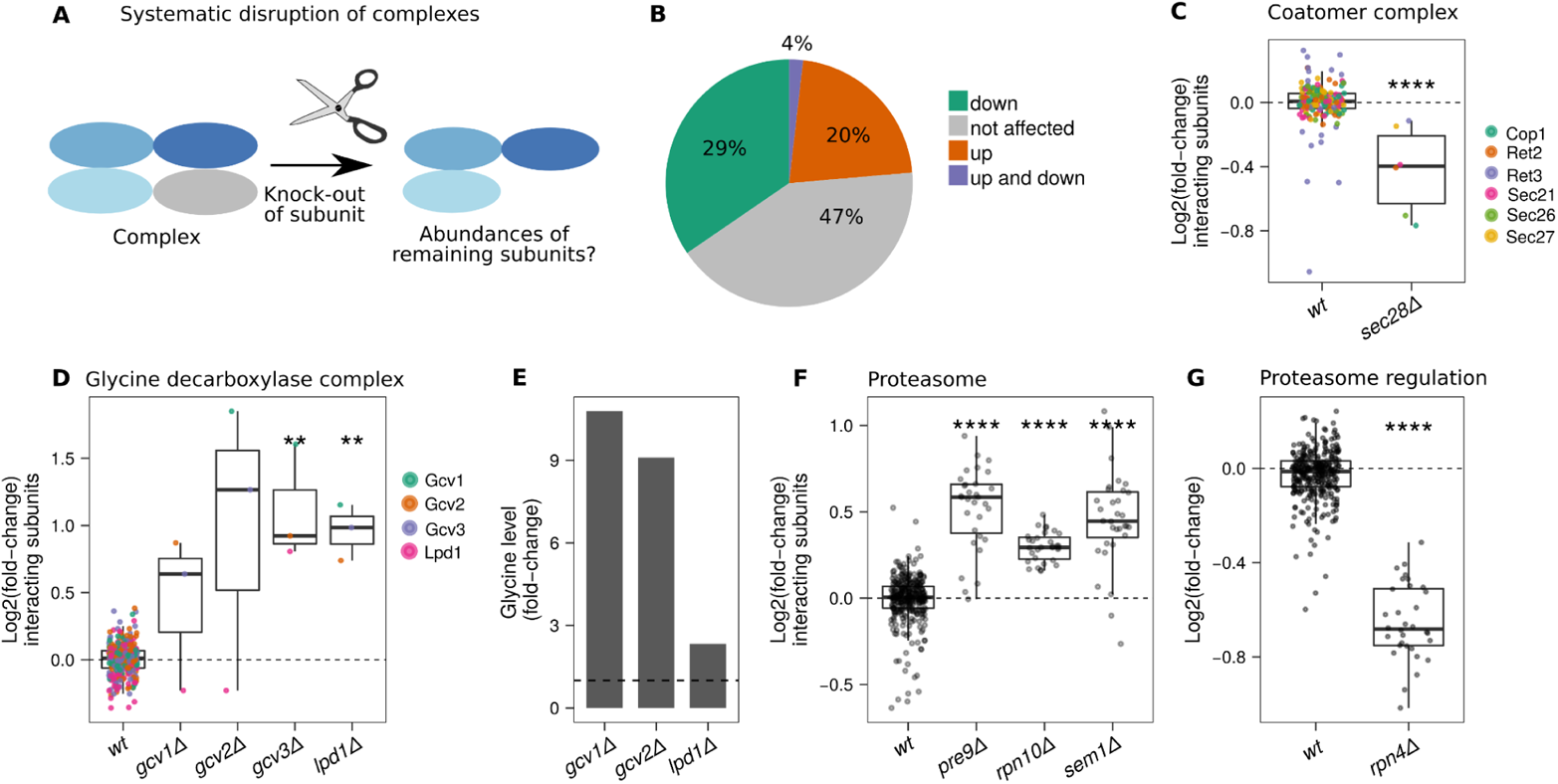
Genome-wide analysis of protein complex perturbation. **(A)** Scheme: Impact a knock-out on the remaining subunits of a protein complex. **(B) The response of protein complexes to perturbation.** Fraction of complexes in which at least one subunit is up-regulated (20%, orange), down-regulated (29%, green), or up- and down-regulated (4%, purple), upon deletion of another subunit. Differential regulation was calculated with a two-sided t-test and by comparing the expression levels of all measured subunits in WT samples with expression levels in the respective knock-outs (adjusted p-value < 0.05, BH for multiple testing correction (Benjamini and Hochberg, 1995)) **(C) The coatomer complex is depleted upon deletion of subunits.** Relative intensities of the subunits Cop1, Ret2, Ret3, Sec21, Sec26, Sec27 are compared between s*ec28Δ* and WT samples. Data is centred and log_2_ transformed. The median expression level across KOs is shown with a dashed line. **(D) Glycine decarboxylase complex, regulated by a metabolic feedback loop, responds to disruption with an up-regulation of other complex subunits.** Relative abundances of the subunits Gcv1, Gcv2, Gcv3 and Lpd1 are shown for the knock-outs of the glycine decarboxylase complex (*gcv1Δ*, *gcv2Δ*, *gcv3Δ*, and *lpd1Δ)* as well as WT samples. Data is centred and log_2_ transformed. **(E) Glycine accumulation in the glycine decarboxylase knock-outs.** Relative glycine concentrations in glycine decarboxylase knock-outs (*gcv1Δ*, *gcv2Δ*, *lpd1Δ*) are shown. The data were centred to the median level across the KOs (dashed line in the plot). A reference dataset was used (Mülleder et al., 2016). **(F) Proteasome complex subunits, regulated by a transcriptional feedback loop, are induced upon deletion of complex members.** The relative protein abundances of all measured proteasome subunits in *pre9Δ*, *rpn10Δ,* and *sem1Δ* (all viable knock-outs of the proteasome complex) are compared to their WT expression levels. Data is centred and log_2_ transformed. The median expression level across KOs is shown with a dashed line. (**G)** Impact of the deletion of the transcription factor Rpn4, which regulates the expression of proteasome complex subunits. The relative protein abundances of all measured proteasome subunits in *rpn4Δ* are compared to their WT expression levels. Significance (two-sided student’s t-test with WT as a reference) is shown with asterisks (**** for p-value ≤ 0.0001; *** for p-value ≤ 0.001; ≤ for p-value ≤ 0.01; * for p-value ≤ 0.05). The first and third quartiles, as well as the median (thick line), are shown with boxplots; whiskers extend to the most extreme data point that is no more than 1.5× the interquartile range from the box.

Notably, we found that 20% of the studied complexes behave differently, and show up-regulation in response to the deletion of at least one subunit (Figure 5B and S5-a). In the search for an explanation, we noted that many complexes in this category are regulated by transcriptional or metabolic feedback loops. For example, the glycine decarboxylase complex, which regulates one-carbon metabolism via methylene tetrahydrofolate (Piper et al., 2000), is up-regulated when glycine levels are high (Sinclair et al., 1996). Indeed, the deletion of a subunit of the glycine decarboxylase complex such as *gcv1Δ* and *gcv2Δ* resulted in an increase in glycine levels (Figure 5E, re-processed data from (Mülleder et al., 2016)) as well as an up-regulation of other complex subunits (Figure 5D). Another example is the proteasome complex (Figure 5F), which is regulated by the short-lived transcription factor Rpn4 via a negative feedback loop to maintain proteasome levels under cellular stress. Indeed, while deletion of the subunits resulted in the upregulation of the other complex members, the deletion of this transcription factor resulted in the down-regulation of the proteasome complex (Figure 5G). Under normal conditions, Rpn4 is degraded by the proteasome. However, upon deletion of proteasome subunits and its resulting reduced function, Rpn4 accumulates and enhances the expression of the proteasome subunits by binding to proteasome-associated control element (PACE) in the promoter region of proteasome subunits (Motosugi and Murata, 2019; Shirozu et al., 2015; Xie and Varshavsky, 2001). Hence, our genome-scale data shines light on the behaviour of protein complex subunits upon their perturbation.

### Large-scale functional proteomics opens new avenues for functionally annotating the proteome

Although several yeast genes have been extensively studied to the degree that they are among the best understood proteins in the literature, many others have received far less attention. For example, the 2,913 best-annotated yeast genes (UniProt annotation score: 5 of 5) have a median of 103 publications each, whereas the 468 worst-characterised genes (score: 1 of 5) are mentioned in a median of 4 publications (Figure S7). We therefore explored the potential of our functional proteomics approach for identifying missing gene functions, reporting four successful and complementary strategies, of which three are specifically enabled by the large-scale combination of functional genomics and proteomics (Figure 6A): i) interpretation of a knock-out strain’s proteome profile (proteomic profiling; PP), ii) interpretation of a protein’s response profile across all other knock-outs (reverse proteomic profiling; RPP), iii) a ‘guilt-by-association’ approach, grouping knock-outs with similar proteome profiles together (profile covariation, or profile similarity), and iv) grouping proteins based on their co-expression across knock-outs (protein covariation).

**Figure 6.**
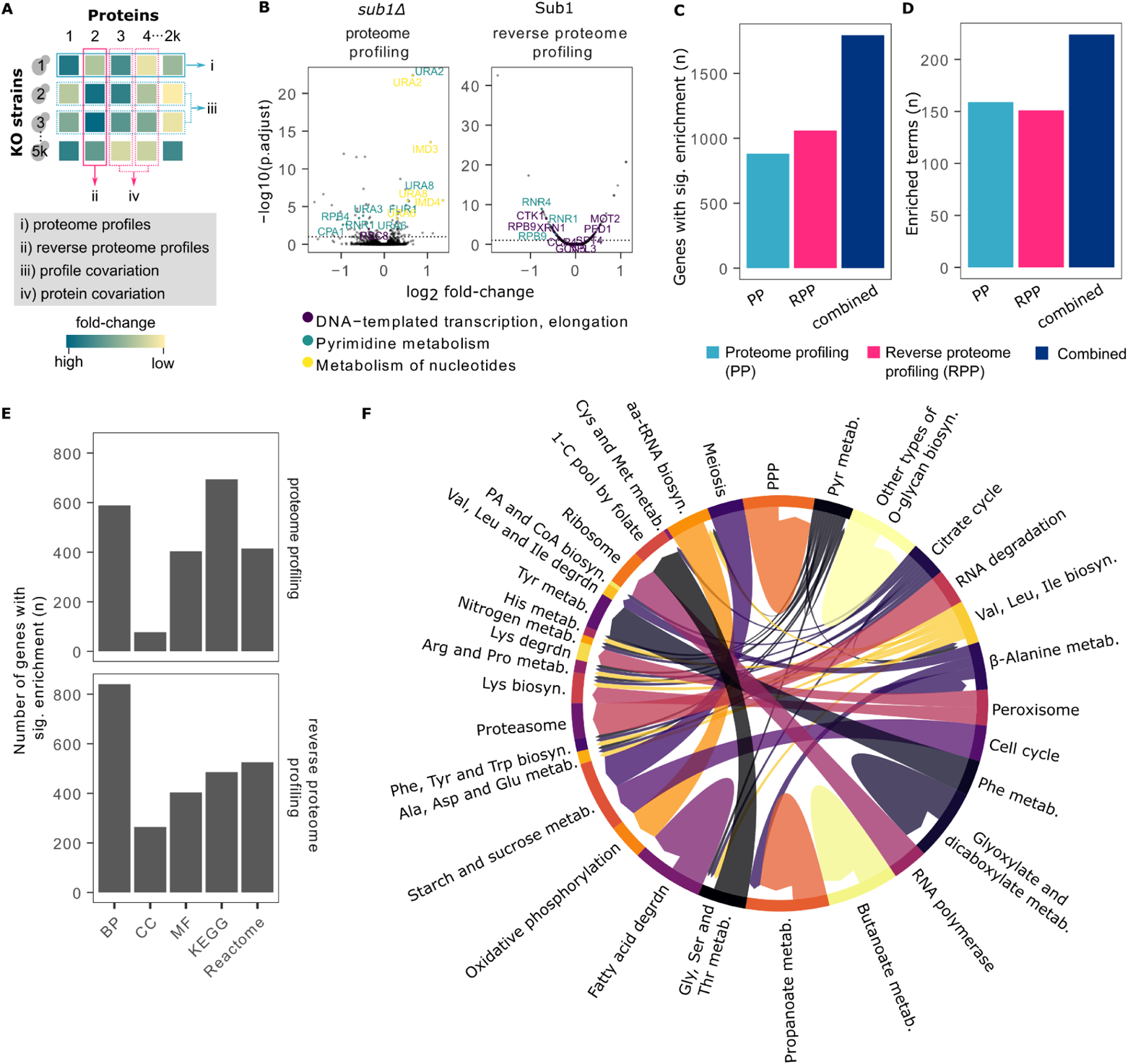
**(A) Genome-scale proteomics facilitates at least four complementary approaches for functional gene annotation. (B) Two complementary functional annotations for the same gene.** Proteome profiling—the response of the proteome in a knock-out (left), and reverse proteome profiling—the response profile of a given protein across knock-outs (right), are shown for the *SUB1* gene as volcano plots. The proteome profiling shows the protein abundance changes of all measured proteins within the *sub1Δ* strain. The reverse proteome profiling shows the protein changes of Sub1 across all KO strains. Centred log_2_ intensities are shown on the x-axes; –log_10_-adjusted p-values are shown on the y-axes. Proteins that were significantly changed (adjusted p-value < 0.01) and that belong to the enriched terms *(DNA-templated transcription, elongation, pyrimidine metabolism, metabolism of nucleotides*) are coloured and labelled. **(C) PP and RPP capture complementary sets of genes.** Number of genes (n) with at least one gene set or pathway significantly enriched by over-representation analysis (p-adjust < 0.05). Numbers are shown for proteome profiling (PP), reverse proteome profiling (RPP) and both profiles combined. GO slim terms (Cherry et al., 2012), KEGG (Kanehisa, 2019; Kanehisa and Goto, 2000), as well as Reactome (Gillespie et al., 2022) were used for the enrichment analysis. **(D) PP and RPP assign complementary functional terms.** Number of terms that were found significantly enriched by over-representation analysis. GO slim terms (Cherry et al., 2012), KEGG (Kanehisa, 2019; Kanehisa and Goto, 2000), as well as Reactome (Gillespie et al., 2022) were used for the enrichment analysis. **(E) Enrichments across gene sets.** Number of genes (n) with at least one term significantly enriched by over-representation analysis (p-adjust < 0.05). Numbers are shown for forward proteome profiling (PP), reverse proteome profiling (RPP), and for both. GO slim terms (Cherry et al., 2012), KEGG (Kanehisa, 2019; Kanehisa and Goto, 2000), as well as Reactome (Gillespie et al., 2022). **(F) Pathway perturbation map for the relationships between perturbations and the response in the proteome.** The KO strains were grouped according to KEGG pathways (Kanehisa, 2019; Kanehisa and Goto, 2000). Arrows face from perturbed pathways to the affected pathways (significantly enriched proteome changes). The latter was assessed by over-representation analysis (hypergeometric test) using the differentially expressed proteins between the KOs and the wild-type strains (limma, p-value < 0.01, BH for multiple testing (Benjamini and Hochberg, 1995)). KEGG terms (Kanehisa, 2019; Kanehisa and Goto, 2000) were used for the over-representation analysis. (Abbreviations: PPP = pentose phosphate pathway; metab. = metabolism; biosyn. = biosynthesis; degrdn = degradation; 1-C = one carbon; PA = pantothenate; aa = aminoacyl; Pyr = pyruvate; Lys = lysine; Phe = phenylalanine; Tyr = tyrosine; Trp = tryptophan; Ala = alanine; Asp = aspartate; Glu = glutamate; Arg = arginine; Pro = proline; His = histidine; Gly = glycine; Ser = serine; The = threonine; Cys = cysteine; Cys = cysteine; Met = methionine; Val = valine; Leu = leucine; Ile = isoleucine).

PP (i) provided functional annotation for 882 genes; in their profiles at least one functional term was enriched (Figure 6C). For example, the knock-out for the RNA polymerase II transcriptional coactivator *SUB1* results in differential expression of many metabolic genes related to pyrimidine and nucleotide metabolism, such as *URA2*, *URA6*, *URA8* (Figure 6B). While PP (i) is a successful conventional strategy to explore gene function and has been extensively used in the past also on knock-out strain collections (Stefely et al., 2016; Zelezniak et al., 2018), it is sensitive to secondary or compensatory mutations (Puddu et al., 2019), and biassed towards non-essential genes that can be efficiently deleted. RPP (ii) instead is complementary both in the sense that high-abundant and often essential proteins are predominantly captured, but also, because it profits from great statistical empowerment, as thousands of proteome profiles are taken into consideration. For example, RPP profiling of *SUB1* functionally associated, in addition to pyrimidine metabolic genes, genes related to *DNA-templated transcription, elongation* such as *CTK1*, *RPB9, PFD1, MOT2* (Figure 6B). Overall, RPP could assign functional terms to 1,060 genes, while both approaches combined (RPP and PP) resulted in an association for 1,794 genes (Figure 6C). The relatively small overlap between PP and RPP (148 genes), as well as different terms assigned, both highlight the complementarity of PP and RPP, respectively. The total number of terms assigned by PP and RPP are 159 and 151, with 73 and 65 being unique to either one or the other profiling approach (Figure 6D and Figure 6E).

Next, we tested whether one could globally map the genetic perturbation to their functional consequences. For this, we grouped the gene-deletion strains on a pathway-by-pathway basis using the KEGG pathway annotation (Kanehisa, 2019; Kanehisa and Goto, 2000). Then, we characterised the proteomic responses to their deletion, also by gene-set analysis (Figure 6F). This global analysis revealed functional relationships between perturbed and responding pathways on a genome scale. Interestingly, we noted that the most common responses to any genetic perturbation were enriched for metabolism, with amino-acid and nucleotide metabolism being among the most frequently responding gene sets (Figure 6G). This result reflects that the metabolic network is a) the largest interconnected biological system in the cell (Ravasz et al., 2002) and b) known to be responsive to general physiological changes (Mülleder et al., 2016). For example, knock-outs related to pyruvate metabolism show proteome responses in various amino-acid metabolic and biosynthetic pathways (i.e. *His, Arg, Pro, Lys, Phe, Tyr, Trp, Ala, Asp, Glu, Gly, Ser, Thr*). We further found that perturbations of the peroxisome result in differential expression in lysine biosynthesis as well as lysine degradation (Figure 5E), reflecting that lysine metabolism is connected to peroxisome deficiency (Breitling et al., 2002). Interestingly, our analysis shows that deleting genes implicated in RNA degradation results in up-regulated proteasomes (Figure 6F, S6A, and S6B), indicating that an increase in RNA levels could be compensated through protein degradation. For example, *mot2Δ* or knock-outs of the LSM complex subunits (*lsm1Δ, lsm6Δ, lsm7Δ)* have increased levels of the proteasome (Figures S6B).

Next, we focussed on associating KO strains with similar proteome profiles (iii), a strategy that was efficient in annotating gene function using both transcriptomics (Kemmeren et al., 2014) and metabolomics (Mülleder et al., 2016). Distance metrics can struggle to calculate meaningful similarities in high-dimensional data as our proteomes (Aggarwal et al., 2001), so we devised a feature-selection strategy based on the observation that proteins which are informative for predicting growth rates are also informative for assessing KO strain similarity (Figures S8, S9; see Methods for details). Selecting 185 (10%) of proteins in this manner strongly improved the detection of functionally related genes (Figure S8E–G). The best similarity metric was found to be robust correlation (Song et al., 2012), and a further performance improvement was achieved by applying a topological overlap measure (Yip and Horvath, 2007). Together, these steps improve the accuracy with which we detect functionally related genes substantially, as shown by a ~30% increase in the area under a PR curve (Figure S10). Finally, we observed that proteome profiles of 2,290 “responsive” KO strains (strains with more differentially expressed proteins than the median strain) could be compared particularly well (Figure S11). We therefore focussed our subsequent analysis of proteome profiles on these responsive strains.

Feature selection also proved beneficial for identifying the second type of associations captured by our data, i.e. those between proteins with similar response patterns across KO strains (Figure 6A). To select the best features for this protein covariation analysis we ranked KO strains by the number of differentially expressed proteins they produced. We found that selecting the 10% most responsive KO strains (467 of 4,675) significantly improved the protein co-regulation analysis (Figure S12). This is approximately the range of features we had previously used for protein covariation analysis (Kustatscher et al., 2019).

### Proteome and KO profiling capture complementary aspects of gene function

Having optimised the computation of proteome profile similarities (of KOs) and the covariation analysis (of proteins), we next asked how well these two approaches capture gene function. For this we performed precision–recall (PR) analyses, testing how well we can identify links between KO strains that are known to be functionally related. We used two different gold standards as reference: functional associations mapped by STRING (Szklarczyk et al., 2019) and interactions between protein-complex subunits mapped by COMPLEAT (Vinayagam et al., 2013). Both proteome profiling and protein covariation detect these associations very well (Figure 7A). To visualise the overall gene–gene (or protein–protein) association dataset we use the dimensionality reduction technique UMAP (McInnes et al., 2018), which creates two maps in which similar KOs (or proteins) are grouped together (Figure 7B). Although our methods do not directly measure physical interactions, grouping proteins by functional similarity means that both maps partially reflect the subcellular organisation of the cell (Figure 7B).

**Figure 7.**
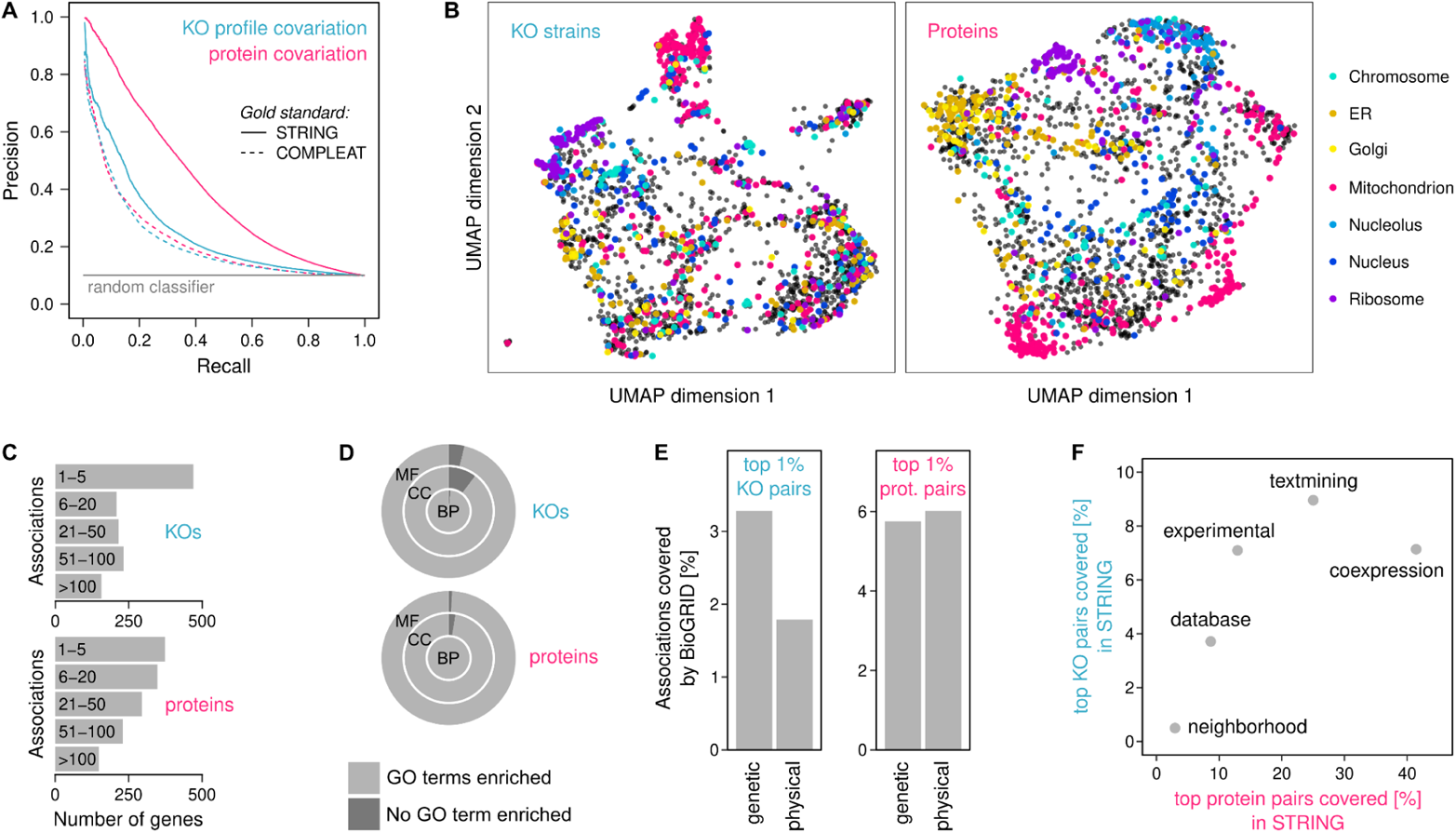
KO profile covariation and protein covariation capture gene function and are highly complementary. **(A)** Precision–recall curves using two different gold standards to show that both profile covariation of knock-outs and protein covariation capture gene function very well. Note this figure focusses on responsive KOs for profile covariation analysis, discarding the non-responsive ones (i.e. gene deletions without differential protein expression). **(B)** UMAPs grouping KO strains by profile covariation (left) and proteins by covariation (right). Subcellular compartments are highlighted and show that both approaches capture subcellular organisation. **(C)** Breakdown of the highest-scoring 1% of associations across each dimension. These cover 1,284 KOs and 1,396 proteins, respectively. **(D)** GO term enrichment analysis for groups of 10 or more connected genes. Nearly all of these are significantly enriched (p < 0.05, Fisher’s exact test) in at least one biological process (BP), and most are enriched in at least one cellular component (CC) and molecular function (MF) term. **(E)** Top 1% of associations were mapped to known interactions in BioGRID, showing that pairs detected by KO profile similarity are more likely to have been previously detected as genetic rather than physical interaction. Co-regulated proteins, on the other hand, are covered better by previously known physical interactions. **(F)** The same associations mapped to known functional associations in STRING and broken down by category. Co-regulated proteins are most similar to (mRNA) co-expression evidence in STRING, whereas proteome profile similarity of knock-outs best reflects associations found by text mining and experimental assays.

We then compared the highest-scoring 1% of pairwise associations found by proteome profile covariation (n = 26,210 KO pairs, Table S6) and protein covariation analysis (n = 26,255 protein pairs, Table S7). They connect a subset of 1,284 KOs and 1,396 proteins, respectively. Some of these genes are linked to fewer than five other genes, others to more than 100 genes (Figure 7C). For both approaches we find that nearly all groups of 10 or more interconnected genes are significantly enriched in at least one GO term (Figure 7D). Nevertheless, there is very little overlap between these top 1% pairwise associations (Figure S13A). This indicates that proteome profiling and KO profiling not only detect different genes (Figure 2A), but also different types of associations between these genes. Indeed, we find that connecting KOs by proteome profile similarity preferentially captures genetic over physical interactions, and associations that were previously detected by literature text mining (Figure 7E,F). In contrast, protein covariation analysis is more likely to capture physical than genetic interactions, and agrees best with associations previously found through mRNA co-expression (Figure 7E,F). Together, these data suggest that proteome and KO profiling provide two highly complementary dimensions for gene-function characterisation. Although protein covariation generally appears to be more precise than proteome profile covariation (Figure 7A, Figure S13B), the best predictive accuracy can be achieved by combining the two scores (Figure S13B).

### Exploring functional relationships within the yeast proteome

To gain more insights into the functional relationships detected by our approach, we explored the behaviour of several example genes in more detail (Figure 8). Dbp3 is an RNA helicase involved in pre-rRNA processing (Weaver et al., 1997), which we cover as knock-out strain and protein. It locates to the nucleolar region of both the KO and protein maps and is linked to other rRNA maturation and ribosome biogenesis factors on both levels (Figure 8A,E). However, the two analyses detect a different subset of ribosome biogenesis factors, reflecting the mentioned complementarity between these two approaches. Similar functional relationships can be explored for all genes that were captured either at KO or protein level (e.g. *SWD3*, Atp14, Arg7) (Figure 8A,E).

**Figure 8.**
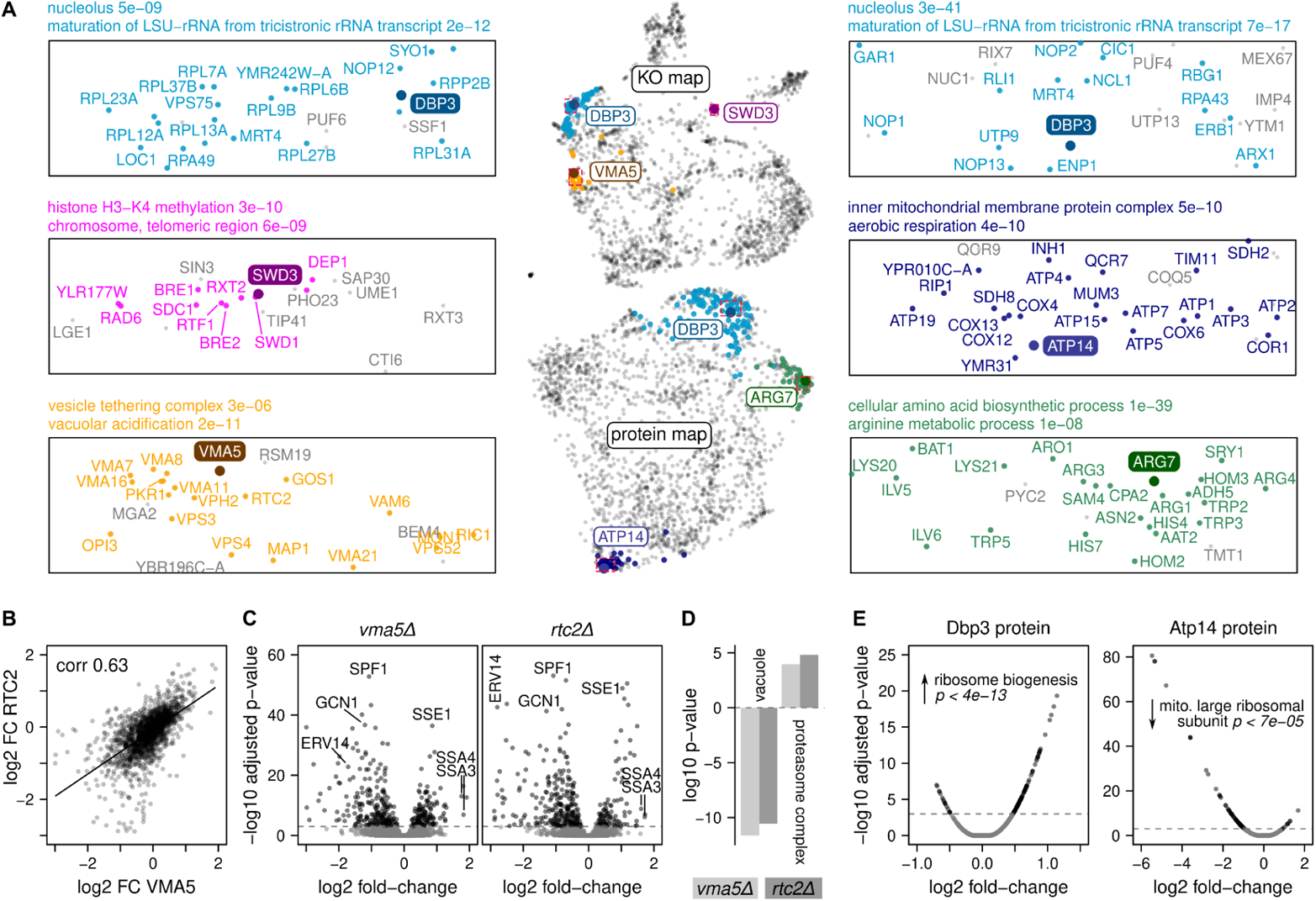
Exploring functional relationships in a proteomic map of genome-scale perturbation. **(A) Six examples showing that proximity in the KO and protein maps reflects functional similarity.** UMAPs of knock-out strains and proteins, with three examples highlighted for each. Subregions that are enlarged and annotated on the left (KOs) and on the right (proteins) are indicated with red dashed lines on the maps. KOs / proteins that are significantly linked to the example gene are highlighted in colour. Significantly linked is defined here as belonging to the 1% highest-scoring pairs in each dataset. Selected GO terms enriched among this group are indicated, together with their enrichment p-value from a Fisher’s exact test. **(B) Proximity in the KO map corresponds to KOs with correlated proteome profiles, suggesting functional similarity.** Protein fold changes (FC) of two KOs that are near each other in the UMAP (*vma5Δ* and *rtc2Δ*, bottom left in (A) are strongly correlated. Correlation coefficient derived from robust correlation (bicor). **(C) The underlying proteome profiles of these KOs provide more detailed insights into which proteins are differentially expressed in each KO.** Many overlapping differentially expressed proteins, a few of which are highlighted, are shown in volcano plots of the proteome profiles of these two KOs. **(D) GO term enrichment of the differentially expressed proteins reveals shared biological trends.** GO term enrichment analysis using a Mann–Whitney U-test shows vacuolar proteins are depleted in both KOs, whereas the proteasome is enriched. **(E) More biological insights about each protein in the protein map can be revealed by analysing their reverse proteome profiles.** Expression of two example proteins, Dbp3 and Atp14 (see also panel (A), across KO strains is shown using volcano plots. Same GO enrichment analysis as in (D), showing that e.g. Dbp3 is up-regulated in KO strains related to “ribosome biogenesis”.

A key advantage of associating KO strains based on their proteome profile, as opposed to simpler phenotypes such as cell fitness, is that proteomes offer detailed insights into how exactly two gene deletions are similar and how they affect cell physiology. For example, the *VMA5* gene encodes a subunit of the vacuolar membrane H^+^-ATPase (Ho et al., 1993). We observe its knock-out to be similar to those of many other genes with vacuolar functions, including genes encoding other H^+^-ATPase subunits (Figure 8A). One of its associated KOs is the putative vacuolar membrane transporter *RTC2*. The proteome profiles of the *vma5Δ* and *rtc2Δ* strains are strongly correlated (Figure 8B) and they share a number of differentially expressed proteins, such as an increase in heat shock proteins Ssa3, Ssa4, and Sse1 (Figure 8C). GO analysis reveals that in both knock-outs vacuolar proteins are down- and the proteasome is up-regulated (Figure 8D). We anticipate that such insights could facilitate hypothesis generation for mechanistic gene-function studies in the future. For example, it is possible that vacuolar defects in the *vma5Δ*, *rtc2Δ*, and related knock-out strains lead to an accumulation of damaged proteins, stimulating the up-regulation of heat shock factors and the proteasome.

## Discussion

A key goal of molecular biology is to gain a deep understanding of biological processes, with a view on explaining phenotypes from the combined impact of genotype and environmental factors. Such genotype–phenotype predictions require an understanding of how biological networks respond to perturbation. Moreover, many genes still lack a functional annotation, and their contribution to the phenotype of an organism is unknown. For example, even though budding yeast is arguably one of the best annotated organism, more than half of the yeast proteome remains understudied (UniProt annotation score below the maximum of 5) and 722 proteins (10%) are classified as functionally uncharacterised by YeastMine (Balakrishnan et al., 2012) (Figure S7). In part, this is due to the “streetlight effect”, a term used to illustrate that current research efforts preferentially focus on a subset of genes that are already relatively well studied (Dunham, 2018; Edwards et al., 2011; Haynes et al., 2018; Sinha et al., 2018; Stoeger and Nunes Amaral, 2020; Stoeger et al., 2018; Wood et al., 2019). There are multiple explanations for this phenomenon: It is easier to work on commonly studied proteins because reagents (antibodies, plasmids, activity assays, mutants, etc.) and important data resources (3D structures, molecular interactions, mutant phenotypes) are more readily available. Moreover, research on widely studied proteins is more likely to be cited in the literature, which can affect visibility of publications, career progression, and funding decisions (Kustatscher et al., 2022).

Genome-scale profiling of loss-of-function mutants has been successfully applied to improve our understanding of biological networks, and delivers gene function as much as they mitigate the streetlighing bias (Giaever et al., 2002; Pan et al., 2004; Tong et al., 2001, 2004). Functional genomic profiling has been extensively applied at the phenotypic level. The Yeast Phenome database (www.yeastphenome.org) collected growth rates of single-gene deletion strains across 5,524 experimental conditions, and the systematic crossing of such strains to create double mutants led to the mapping of 900,000 genetic interactions (Costanzo et al., 2010, 2016). Our study provides a significant amount of data to help interpret the phenotypic output of these screens at the molecular level. Moreover, the proteome is complementary, and provides added value, to other ‘functional omic’ screens, including genome-scale transcriptomics (Kemmeren et al., 2014) or metabolomics (Mülleder et al., 2016) studies that do not capture the post-transcriptional regulation of protein function. For instance, we herein identify a number of protein complexes for which degradation of surplus subunits is induced when the complex is disrupted. Equally though, by showing the mechanism is not universal, we provide a differentiated picture about protein complex dynamics. Indeed, we discover that 20% of protein complexes have subunits induced upon perturbations. Our dataset hence allows us to systematically study post-transcriptional processes that have previously been studied in isolation, or that are detected neither by phenotypic screens, transcriptomics or metabolomics, respectively, and interpret the findings quantitatively.

With the genome-scale size of the proteomic dataset, and due to the link of each perturbation to a specific gene, one identifies properties that are of general importance to explain protein expression. We describe an interdependency of protein abundance with protein half-life and ribosome occupancy, suggesting these could be general drivers of differential protein expression. Moreover, we observe that the direction of protein expression changes is restrained in a protein-specific manner; most proteins are preferentially down-regulated or preferentially up-regulated, while only a few proteins change in either direction. Protein turnover is partially linked to gene essentiality, with essential genes generally having faster turnover rates (Figure S4D). Our data hence establish that protein expression dynamics and turnover are interlinked and not independent properties. Consequently, we provide a differentiated picture of protein expression in general. On the one hand, we confirm and quantify the paradigm that differential protein expression is driven by function. Paralogues and proteins connected in genetic, metabolic, evolutionary, or protein–protein interaction networks have a higher likelihood to respond to deletion. Equally, however, the general biophysical and biological properties of the proteome, like the location of a protein on a potential aneuploid chromosome, or its turnover rate, have a strong contribution to the response of the proteome to perturbation as well. Our study provides the community with data to facilitate the distinction of function-specific and function-independent differential protein expression, to map the responsible regulatory networks.

Our study shows that genome-scale genetic perturbation and proteomics are a powerful combination for annotating, or improving the annotation, of gene function. On the one hand this is due to the large-scale nature of our investigation: in contrast to analysing the proteome of an individual knock-out strain, our approach benefits from three additional dimensions that are made possible through comparing the profiles with each other, or by studying coexpression patterns. These are useful for studying gene function, both, because they are complementary to the interpretation of an individual profile, as well as profit from the statistical empowerment of a massive scale dataset. The new approaches introduced with our study include what we coin a “reverse proteome profile”, i.e. identifying the type of gene deletions that trigger an expression change of a particular protein. It also includes two separate guilt-by-association approaches (Marcotte et al., 1999; Vazquez et al., 2003), which infer gene function by associating genes with similar knock-out proteome profiles and proteins with similar expression patterns, respectively. These approaches are attractive not only because of their complementary nature, but also because they exploit the statistical possibilities provided by the dataset to mitigate confounding factors that exist within each genetic screen. Indeed, the widespread use of the KO collection has revealed some shortcomings that apply to all genetic-perturbation-based approaches. One of them is that eukaryotic genomes are versatile and compensatory, with secondary genetic adaptations readily appearing. For instance, any mutation that results in mitochondrial genome loss, because it affects mitochondrial respiration, the mitochondrial ribosome, or iron metabolism, selects for petite-suppressing mutations in *ATP3* or *ATP1*, respectively, that restore the mitochondrial membrane potential in the absence of a mitochondrial genome (van Leeuwen et al., 2016; Vowinckel et al., 2021). Another example are aneuploidies, that can emerge as a transient adaptation to stress (Yona et al., 2012). However, errors resulting from such general confounders in gene function annotation can be mitigated by comparing profiles across many strains, or by studying protein covariation. Moreover, such confounders can more easily be identified with a large yet precise dataset at hand. For instance, we identify the gene-knockouts resulting in chromosomal aneuploidies, and find that these explain broad proteomic profiles known to be frequent in slow-growing strains.

Finally, part of the power of the approach is provided by the complementary nature of genetic perturbations and proteomics. Proteomics is helpful for studying essential genes, which tend to produce abundant proteins but cannot be deleted, while gene deletion works well for non-essential and low-abundant proteins which cannot be detected by MS. Hence, the combination of the two technologies allowed us to identify far more functional associations between yeast genes than what could have been achieved using either technology alone. Similar synergistic findings were reported by a recent study integrating immunofluorescence images with protein-complex affinity-purification data (Qin et al., 2021). Thus, the combination of multiple large-scale technologies with complementary strengths and biases could become a paradigm for providing accurate and comprehensive data-driven gene-function annotation. This is especially relevant for future studies addressing the problem of understudied proteins. Indeed, the problem of understudied genes is more pronounced in higher organisms, and much more in all non-model organisms (Kustatscher et al., 2022; Warren, 2015). The approaches developed as part of our paper can be applied to any kind of large-scale proteome, including mammalian ones, where through the development of CRISPR/Cas9 genetic libraries have become increasingly available (Peng et al., 2015).

## Acknowledgments

We thank R. King, R. Lane, E. Hudson, N. Morrice, P. Brooks and J. B. Vincendet for their help with TripleTOF 6600. We thank Michael Howell and the Crick HTP for helping in filling the culture plates. We thank Juri Rappsilber for discussions about data analysis strategies. This work was supported by the BBSRC (nos. BB/N015215/1 and BB/N015282/1), the Francis Crick Institute, which receives its core funding from Cancer Research UK (no. FC001134), the UK Medical Research Council (no. FC001134) and the Wellcome Trust (no. FC001134 and IA 200829/Z/16/Z), the European Research Council (ERC) under grant agreement ERC-SyG-2020 951475, as well as the Ministry of Education and Research (BMBF), as part of the National Research Node ‘Mass spectrometry in Systems Medicine (MSCoresys), under grant agreement 031L0220. GK is funded by an MRC Career Development Fellowship (MR/T03050X/1).

## Data availability

The processed quantitative data files have been uploaded to Mendeley data (https://data.mendeley.com/datasets/mk6hw2zt7m/draft?a=73ec76b8-3e4c-4c88-b00b-96529ecc21a6). The gene networks were downloaded from Yeastnet (Kim et al., 2014). The complex data was downloaded from EBI Complex Portal (Meldal and Orchard, 2018; Meldal et al., 2015, 2019). The ribosome profiling data (McManus et al., 2014), protein turnover rates (Martin-Perez and Villén, 2017) and glycine concentrations (Mülleder et al., 2016) were downloaded from the respective reference datasets. GO slim terms were downloaded from the Saccharomyces Genome Database (Cherry et al., 2012).

## Materials and Methods

### Materials

Yeast nitrogen base without amino acids (Sigma, Y0262), D-(+)-Glucose (Sigma, G7021), water (LC-MS Grade, Optima; 10509404), acetonitrile (LC-MS Grade, Optima; 10001334), formic acid (LC-MS Grade, Thermo Scientific Pierce; 13454279), and methanol (LC-MS grade, Optima, A456-212) were purchased from Fisher Chemicals. DL-dithiothreitol (BioUltra, 43815), iodoacetamide (BioUltra, I1149), solid-glass beads (borosilicate, diameter 4 mm, Z143936-1EA), and ammonium bicarbonate (eluent additive for LC-MS, 40867) were purchased from Sigma Aldrich. Urea (puriss. P.a., reag. Ph. Eur., 33247H) and acetic acid (eluent additive for LC-MS, 49199) were purchased from Honeywell Research Chemicals. Trypsin (Sequence grade, V511X) was purchased from Promega. iRT peptides (Ki-30002-b) were purchased from Biognosys. MS Synthetic Peptide Calibration Kit (5045759) was purchased from SCIEX.

### Yeast culture

The *Saccharomyces cerevisiae (*S228c) haploid (MATa) deletion collection (Winzeler et al., 1999) with restored prototrophy (Mülleder et al., 2012) was used. The single knock-out strains were arranged on 96-well plates. Each plate, except for plate 17, contained 7 wild-type strains in positions A11, B8, C5, D2, F11, G8, and H5. Plate 17 contained 5 wild-type strains in positions B8, C5, D2, F11, G8, and H5.

The yeast strains were grown in batches of 12 96-well plates. In order to reduce batch effects, the media for all batches was prepared at once, pre-filled into 96-well plates and stored at –80°C until the day of the experiment. Further, a 5x SM medium stock solution was prepared and stored at –80°C and used for the agar plates, which were prepared fresh on the day of the experiment. All media were filtered (0.22 µm filter, GP Millipore Express Plus membrane) and the plates as well as the beads were autoclaved before usage. All pipetting was done with a liquid-handling robot (Biomek NX^P^ automated liquid handler) and yeast cells were pinned with a pinning robot (Rotor, Singer instruments).

The yeast strains were grown as previously published (Mülleder et al., 2016) with slight modifications. The thawed stock cultures were spotted with the pinning robot onto synthetic minimal (SM) agar medium (6.7 g/l yeast nitrogen base without amino acids, 2% glucose, 2% agar) and incubated at 30°C for 47–49 hours. Subsequently, these cells were used for inoculation in 200 μl SM liquid medium in 96-well plates and incubated at 30°C. After 19.75 hours, 160 μl culture was transferred to a deep-well plate (ABgene storage plates, AB-0661) prefilled with 1,440 μl SM liquid medium (1/10 dilution) and with one solid-glass bead (borosilicate) per well. The plates were sealed with a membrane (Breathe-Easy sealing membrane for multiwell plates, Diversified Biotech, Z763624-100EA) and incubated for 8 hours at 30°C with 1,000 rpm mixing (Heidolph Titramax incubator). Subsequently, the culture was transferred into a fresh 96-well plate (Eppendorf, 10052143) and spun down at 4,000 rpm (Eppendorf Centrifuge 5810R). The supernatant was removed and the plate was sealed with aluminium foil (adhesive PCR plate foil, Thermo Scientific, AB0626) as well as a plastic lid (CLS3098) before being frozen and stored at –80°C until further processing.

### Sample preparation

The protein extraction and digestion was done in batches of 4 plates (384 samples). In order to reduce batch effects, stock solutions (120 mM iodoacetamide, 55 mM DL-dithiothreitol, 9 μl 0.1 mg/ml trypsin, 2 μl 4x iRT) were prepared at once and stored at –80°C. Other stock solutions (7 M urea, 0.1 M ammonium bicarbonate, 10% formic acid) were stored at 4°C. All pipetting was done with a liquid-handling robot (Biomek NX^P^ automated liquid handler), shaking was done with a thermomixer (Eppendorf Thermomixer C) after each step, and for incubation a Memmert IPP55 incubator was used.

200 μl 7 M urea / 100 mM ammonium bicarbonate and glass beads (~100 mg/well, 425–600 µm, Sigma, G8772) were added to the frozen pellet. Subsequently, the plates were sealed (Cap mats, Spex, 2201) and lysed using a bead beater for 5 min at 1,500 rpm (Spex Geno/Grinder). After 1-min centrifugation at 4,000 rpm, 20 μl 55 mM DL-dithiothreitol were added (final concentration 5 mM), mixed, and the samples were incubated for 1 h at 30°C. Subsequently, 20 μl 120 mM iodoacetamide were added (final concentration 10 mM) and incubated for 30 min in the dark at room temperature. 1 ml 100 mM ammonium bicarbonate was added, centrifuged for 3 min at 4,000 rpm, then 230 μl were transferred to prefilled trypsin plates. After incubation of the samples for 17 h at 37°C, 24 μl 10% formic acid were added. The digestion mixtures were cleaned up using C18 96-well plates (96-Well MACROSpin C18, 50–450 μL, The Nest Group, SNS SS18VL). For solid-phase extraction, 1 min of centrifugation at the described speeds (Eppendorf Centrifuge 5810R) was used to push the liquids through the stationary phase and the liquid handler was used to pipette the liquids onto the material. The plates were conditioned with methanol (200 μl, centrifuged at 50 *g*), washed twice with 50% ACN (200 μl, centrifuged at 50 *g*, then the flow-through discarded), equilibrated three times with 3% ACN, 0.1% FA (200 μl, centrifuged at 50, 80, 100 *g*, respectively, then the flow-through discarded). 200 μl of digested samples were then loaded (centrifuged at 100 *g*) and washed three times with 3% ACN, 0.1% FA (200 μl, centrifuged at 100 *g*). After the last washing step, the plates were centrifuged another time at 180 *g* before the peptides were eluted in 3 steps (twice with 120 μl and once with 130 μl) 50% ACN (180 *g*) into a collection plate (1.1 ml, square well, V-bottom). Collected material was completely dried in a vacuum concentrator (Eppendorf Concentrator Plus) and redissolved in 40 μl 3% ACN, 0.1% formic acid before being transferred into a 96-well plate (700 μl round, Waters, 186005837) prefilled with iRT peptides (2 μl, 4x diluted). QC samples for repeat injections were prepared by pooling digested and cleaned-up samples from 4 different 96-well plates.

2 μl of each sample were loaded onto the microfluidic 96 Lunatic plates. Peptide concentrations were measured with the Lunatic instrument (Unchained Labs). Protein concentrations were calculated from the absorbance value at 280 nm and the protein-specific extinction coefficient.

### Liquid chromatography–mass spectrometry

The digested peptides were analysed on a nanoAcquity (Waters) (running as 5 µl/min microflow LC) coupled to a TripleTOF 6600 (SCIEX). 2 µg of the yeast digest (injection volume was adjusted for each sample based on the measured peptide concentration) were injected and the peptides were separated in a 19-min nonlinear gradient (Table S2) ramping from 3% B to 40% B (Solvent A: 1% acetonitrile/0.1% formic acid; solvent B: acetonitrile/0.1% formic acid). A Waters HSS T3 column (150 mm x 300 µm, 1.8 µm particles) was used with a column temperature of 35°C. The DIA method consisted of an MS1 scan from m/z 400 to 1250 (50 ms accumulation time) and 40 MS2 scans (35 ms accumulation time) with variable precursor isolation width covering the mass range from m/z 400 to 1250 (Table S1). Rolling collision energy (default slope and intercept) with a collision energy spread of 15 V was used. A DuoSpray Ion Source was used with ion source gas 1 (nebuliser gas), ion source gas 2 (heater gas), and curtain gas set to 15 psi, 20 psi, and 25 psi. The source temperature was set to 0°C and the ion-spray voltage to 5,500 V. The measurements were conducted within a period of 12 month and on 2 different platforms with identical setups (Figures S1b).

### Growth assays

Growth assays were performed on SC, SM and YPD by time-course imaging of colonies, using our Pyphe pipeline (Kamrad et al., 2020), as described in (Kamrad et al., 2022). Library plates were grown from cryostocks in 384 format for three days on agar media. Plates were then multiplexed into 1,536 format on agar with two grids of 96 wild-type controls (BY4741 DeltaHIS3 pHLUM) placed in the top-left and bottom-right corners. Plates were then passaged again and copied onto fresh agar plates which were immediately placed into a transmission scanner (Epson V800) located in an incubator maintained at 30°C. Plates were imaged approximately every 20 min for 40 h. Growth curves based on pixel intensity values were extracted and smoothed using a median and Gaussian filter with kernel sizes of 3. Maximum slopes were then extracted using a sliding window of length 5. Grid values in the bottom-left and top-right corner were extrapolated using linear regression. Maximum slopes were normalised by grid correction (Zackrisson et al., 2016, G3) and repeats for the same knock-out were averaged. Assay plates consistently exhibited signal-to-noise ratios above 30 and fractions of unexplained variance below 20%, indicating high data quality.

### Library generation

The libraries were generated from “gas-phase fractionation” runs using scanning SWATH and small precursor isolation windows. 5 µg yeast digests were injected and run on a nanoAcquity UPLC (Waters) coupled to a SCIEX TripleTOF 6600 with a DuoSpray Turbo V source. The peptides were separated on a Waters HSS T3 column (150 mm x 30 0µm, 1.8 µm particles) with a column temperature of 35°C and a flow rate of 5 µl/min. A 55-min linear gradient ramping from 3% ACN/0.1FA to 40% ACN/0.1% FA was applied. The ion source gas 1 (nebuliser gas), ion source gas 2 (heater gas), and curtain gas were set to 15, 20, and 25. The source temperature was set to 75 and the ion spray voltage to 5,500 V. In total 11 injections were run with the following mass ranges: m/z 400–450, 445–500, 495–550, 545–600, 595–650, 645–700, 695–750, 745–800, 795–850, 845–900, 895–1000, and 995–1200. The precursor isolation window was set to m/z 1 except for mass ranges m/z 895–1000 and 995–1200, where the precursor windows were set to m/z 2 and 3, respectively. The cycle time was 3 sec consisting of high and low energy scan and data was acquired in “high resolution” mode. A spectral library was generated using library-free analysis with DIA-NN directly from these scanning SWATH acquisitions. The UniProt (The UniProt Consortium, 2017) yeast canonical proteome was used for library annotation.

### Raw data processing

Raw data processing was carried out with DIA-NN (Version 1.7.12) with default settings with MS2 and MS1 mass accuracies set to 20 ppm and scan window size set to 6. Precursors were filtered for q-values < 0.01 (precursor and protein level) and only proteotypic peptides were considered. Batches (plates) were corrected by bringing median precursor quantities of each batch to the same value (dividing the quantities by the plate median and multiplying them with the median of all plate medians). Precursors were only considered if identified in > 80% of WT samples and if quantified with CV < 50%. Samples were removed if the number of identified precursors was less than 80% of the maximum number of precursors. Protein quantities were obtained using the MaxLFQ algorithm (Cox et al., 2014) as implemented in the DIA-NN R package (https://github.com/vdemichev/diann-rpackage). Missing values were imputed with a mixed imputation strategy. Protein quantities that were missing in < 5% of the samples per plate, were imputed with a random value between 0 and the minimum protein quantity per plate. Values that were missing in > 5% of the samples per plate were imputed with nearest neighbour averaging (KNN) using the impute.knn function from the R package impute (Hastie et al., 1999; Troyanskaya et al., 2001). Plates 59 and 60 were excluded from further analysis as they contained mainly WT samples. Plate 54 was excluded as the position that is supposed to contain a blank sample (unique position for each plate) contained a yeast sample, indicating that the plate was most likely wrongly positioned in the autosampler.

### Data analysis

All statistical analyses were done in R (v.3.6.3) (RDc). For basic data manipulation and visualisation the R tidyverse group of packages were used (Wickham et al., 2019).

Differential expression analysis on the log_2_-transformed data was conducted with the limma R package (Ritchie et al., 2015). Genotypes (or functional groups) were included in the linear model while only the genotypes (or functional groups) were defined in the contrast matrix. Further, an intensity trend in the prior variance was allowed in the empirical Bayes statistics. Coefficients of variation were calculated for each protein or precursor as its empirical standard deviation divided by its empirical mean, and are reported in percentages. CV values were calculated for proteins or precursors identified in at least two replicate measurements. Heatmaps were generated with the ComplexHeatmap R package and default settings (Gu et al., 2016). Enrichment analyses (hypergeometric tests) were performed with the piano package(Väremo et al., 2013) using all knock-outs as background.

Centered protein intensities were calculated by dividing the protein intensities by the median of all knock-out and WT samples. *Z*-scores were calculated by dividing the (centred) protein quantities by their standard deviations. The measurements of the individual knock-out strains were not replicated. No statistical method was used to calculate sample size prior to the experiment.

For the generalised linear models with elastic net, the glmnet function within the caret package (Kuhn, 2008) was used. A tune length of 5 with 10-fold cross-validation was applied to generate the models. The data was scaled, centred, and split into train and test dataset (80 /20). Feature selection was done with varimp function within the caret package (Kuhn, 2008).

Complexes with less than 3 measured proteins were excluded in the analysis. In addition, the following complexes were removed before the analysis due to redundancy in subunits: CPX-1882, CPX-1883, CPX-776, CPX-1675, CPX-473, CPX-1602, CPX-769, CPX-770, CPX-771, CPX-776, CPX-581, CPX-44, CPX-32, CPX-1102.

### Data transformation for KO and proteome profiling

For proteome profile similarity assessment of KO strains, protein intensities were divided by the median intensity across all strains (WT, KO, and QC samples) and log_2_-transformed. The resulting data matrix contained relative protein level changes of 1,850 proteins across 5,463 samples without missing values (see above for imputation strategy). For protein covariation analysis, protein intensities were transformed in the same way but starting from a non-imputed and less stringently filtered data matrix (considering precursors identified in > 50% rather than 80% of WT samples), because this type of analysis is not affected by a moderate amount of missing values (Kustatscher et al., 2019). The resulting data matrix contained 2,292 proteins across 5,552 samples with 7.5% missing values.

### Profile comparisons using correlation and distance metrics

To avoid spurious correlations between proteome profiles, log_2_ fold changes were normalised such that the median fold change of each protein across KOs was zero. To avoid spurious correlations between KO profiles, log_2_ fold changes were normalised such that the median protein fold change of each KO was zero. We tested a range of similarity metrics, including three correlation metrics, three “conventional” distance metrics (Euclidean, Manhattan, Minkowski), and two decision-tree-based distance metrics. Input data were scaled (z-transformed) prior to calculation of conventional distance metrics. Pearson and Spearman correlations, as well as Euclidean, Manhattan, and Minkowski distances were calculated using base R functions. Biweight midcorrelation (bicor) was applied through the WGCNA R package (Langfelder and Horvath, 2008, 2012). The treeClust R package (Buttrey and Whitaker, 2015) was used to calculate distances with the treeClust algorithm, using default parameters except for *minsplit* = 500, which had been identified as the optimal parameter setting using precision–recall test runs. Unsupervised random forests (uRFs) were used through the randomForest R package (Liaw and Wiener, 2002). Note that uRFs do not work on datasets with missing values, so for covariation analysis via uRFs missing values were imputed using the *k-*nearest-neighbour imputation algorithm of the impute R package (Hastie et al., 2021).

The topological overlap matrix was calculated using the TOMsimilarity function of the WGCNA R package (Langfelder and Horvath, 2008; Zhang and Horvath, 2005).

### Precision–recall analysis

Precision–recall (PR) curves and the areas under these curves were calculated using the PRROC R package (Grau et al., 2015). Modified PR curves, in which recall is shown as the number of retrieved true positives at log_10_ scale, were created using the ROCR R package (Sing et al., 2005).

We used two separate, partially overlapping gold standards for the PR analyses in this study: one based on functional protein–protein associations reported by STRING v11 (Szklarczyk et al., 2019) and one based on the COMPLEAT set of protein complexes (Vinayagam et al., 2013). For the STRING gold standard, true positive (TP) associations were defined as gene pairs with a combined STRING score of ≥ 700 (high confidence). False positive (FP) pairs were defined as all pairs that were not linked by STRING at any confidence level. The COMPLEAT gold standard was described previously (Mülleder et al., 2016; Vinayagam et al., 2013). From both gold standards we further excluded FP pairs that had been found associated by either STRING, COMPLEAT, BioGRID v3.5 (Stark et al., 2006) or Gene Ontology (Myers et al., 2006). In addition, we removed all genes that had not been detected as part of the Y5K dataset, and those that could not be unambiguously cross-mapped between UniProt IDs and systematic gene names (OLNs). The resulting gold standards contain 70,023 unique STRING TPs, 58,785 unique COMPLEAT TPs, and 14,726 TPs that overlap between the two standards.

### Feature selection for gene function prediction

Selecting random subsets of the available 1,850 proteins indicated that our analysis could be improved by feature selection (Figure S8A). We therefore aimed to systematically select the best features (i.e. proteins) to link KO strains. In principle, it would be possible to identify the optimal subset of features for this task simply by selecting those that result in the largest area under the PR curve. However, such a “cherry-picking” approach may not extrapolate well to other data sets or gold standards. We therefore based our feature selection process on the prediction of growth rates. Our rationale was that proteins which are important for growth rate prediction may also be the ones whose expression changes are relevant for linking KO strains (see also legends of Figure S8 and S9 for additional explanations).

For this feature selection process we took advantage of the ability of random forests (RFs) to determine the importance of individual features (i.e. proteins) for a regression task (Breiman, 2001). We used the randomForest R package (Liaw and Wiener, 2002) to train RF regression models on the growth rates of all KO and wildtype strains. We trained three separate RFs (technical replicates) for each of the three growth media (SC, SM, YPD) for which growth rates had been measured. These 9 RF models were created using default parameters except for *nodesize*, which was set to 100 to speed up the calculation. To test if RF regression models can accurately predict growth rates, we created a 10th model in which we withheld 500 strains from the training set and predicted their growth rates in YPD medium (Figure S3B).

Feature importance was determined as the increase in node purity for each protein (“IncNodePurity” output from the RF models). Under the chosen parameter settings we found this measure of feature importance to be highly reproducible between technical replicates, i.e. RF models trained on the same input data (R^2^ = 0.99). However, feature importance differed considerably for growth rate predictions in the three growth media (e.g. R^2^ = 0.65 between SM and SC). Feature importances from different RF models were scaled (z-transformed) and proteins were ranked by the minimum importance they achieved across different RF models.

To select the best features (KO strains) for protein covariation analysis, KO strains were ranked by the number of differentially expressed proteins in decreasing order. The most responsive 10% of KO strains selected in this way proved to be the ideal set of KO strains to use for protein covariation analysis (Figure S12).

### UMAP visualisation

The R implementation of the UMAP algorithm (Konopka, 2020; McInnes et al., 2018, 2020) was used to reduce protein and KO correlation matrices down to two dimensions. Since UMAP uses distances and not similarities to calculate the low dimensional projection of the data, biweight midcorrelations were inverted (multiplied by –1) before UMAP analysis.

### Additional data annotation

For the gene categorisation in Figure 2A, essential yeast genes were defined as those annotated as “inviable” in the Saccharomyces Genome Database (Cherry, 2015). A list of uncharacterised yeast genes was downloaded from YeastMine (Balakrishnan et al., 2012). Protein lengths were extracted from UniProt (The UniProt Consortium, 2017). Protein abundances for Figure 2A, which had to cover proteins that were not detected in this analysis, were extracted from a meta-analysis of absolute protein concentrations in yeast (Ho et al., 2018). Gene Ontology (GO) term enrichment for the vma5Δ and rtc2Δ strains (Figure 8) and the 185 feature-selected proteins (Figure S9A) were carried out using the Panther website as described (Mi et al., 2013). Large-scale GO enrichment analysis as described in Figure 7D was performed using the topGO R package (Alexa and Rahnenfuhrer, 2016).

