## Supplementary information for "The Proteomic Landscape of Genome-Wide Genetic Perturbations"

**Index**


| | |
|---|---|
| Page 3–22 | Supplementary Figures S1–S13 |
| Page 23–28 | Supplementary Tables S1–S5 |
| Page 29 | Supplementary References |

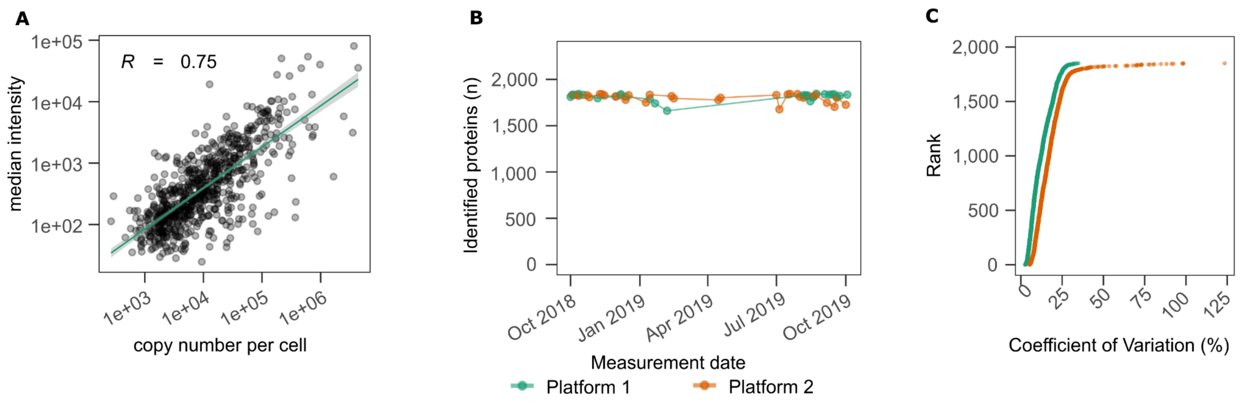


**Figure S1.** (**A**) Median intensity values across all WT samples are plotted against copy numbers per cell taken from a reference dataset (Lawless et al., 2016). Scales are $\log_{10}$ transformed. (**B**) The identification numbers are consistent and independent of time and instrument used. The data were acquired within a time frame of 12 month. (**C**) The coefficients of variation (in %) were calculated for whole-process control samples (green, n = 375), and KO samples (orange, n = 4,703).

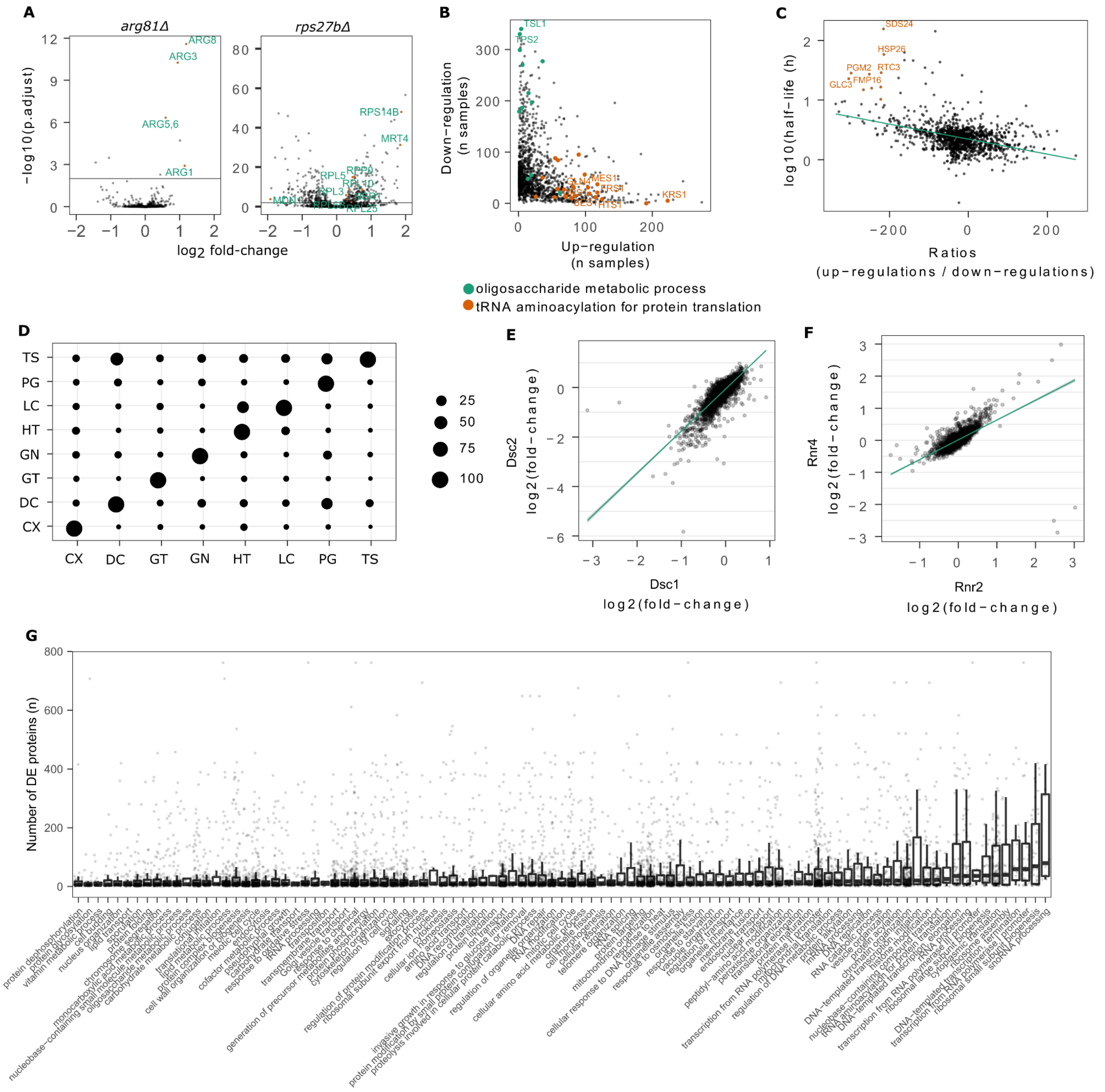


**Figure S2. (A)** Differential protein expression in *arg81Δ* (left) and *rps27bΔ* (right) strains. While *arg81Δ* has a specific proteome response with a low number of differentially expressed proteins, many proteins are affected in *rps27bΔ*. Differential expression was calculated with the limma package (Ritchie et al., 2015) and BH was used for multiple testing (Benjamini and Hochberg, 1995). The x-axis shows centred $\log_2$ intensities and the y-axis shows adjusted p-values ($-\log_{10}$ transformed). **(B)** Up-regulations (x-axis) and down-regulation (y-axis) of each protein across the 4,567 KO strains. Proteins related to oligosaccharide metabolic process (green) and tRNA aminoacylation for protein translation (orange) are labelled. **(C)** The directionality of differential expression is given as ratios (number of up-regulations / number of down-regulations) for each protein and is plotted against its protein half-life (y-axis). Protein half-lives were $\log_{10}$ transformed. **(D)** Overlap between genetic, physical, and functional interaction networks. The overlap is normalized to the size of the network given on the y-axis. Numbers are given in %, with 100 indicating a complete overlap. Interactions were downloaded from YestNet (v3, (Kim et al., 2014)) (LC = literature curated PPI; TS = tertiary structure of protein; HT = high-throughput PPI; GN = genomic neighbor; CX = co-expression; GT = genetic interaction; DC = domain co-occurrence; PG = phylogenetic profiles). **(E,F)** Examples of co-expressed paralogues. Centred protein intensities are plotted against each other for Dsc2 / Dsc1 and Rnr4 / Rnr2. **(G)** Number of differential expressions for each gene deletion, grouped by Gene Ontology slim terms for *biological process (Cherry et al., 2012)*. Differential expression was calculated with the limma package (Ritchie et al., 2015) and BH was used for multiple testing (Benjamini and Hochberg, 1995).

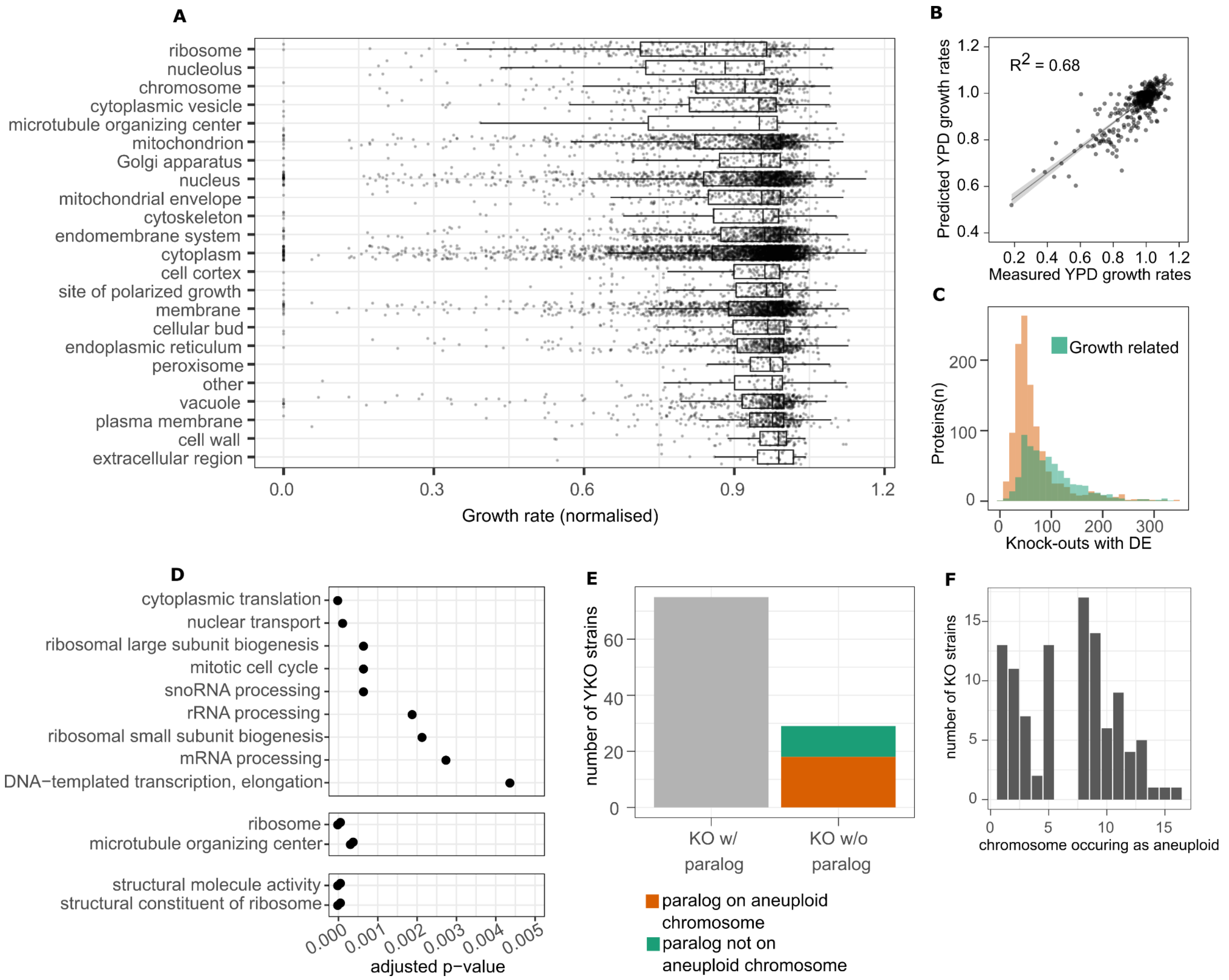


**Figure S3. (A)** Growth rates (normalised) for each knock-out strain, grouped by cellular compartments (GO slim terms for *cellular compartment* (Cherry et al., 2012)). The first and third quartiles, as well as the median, are shown with boxplots, and the whiskers extend to the most extreme data point that is no more than 1.5× the interquartile range from the box. (**B**) Growth rates were predicted from the protein abundances using a random forest (RF) algorithm. Growth rates in YPD medium were measured for all strains. We then trained an RF regression model to predict these KO strain growth rates from the abundances of the 1,850 quantified proteins. 500 KO strains were left out from training the RF regression model, which was subsequently used to predict their growth rates in YPD medium. See also Figure S8 and Methods. (**C**) The proteomic changes in knock-out strains are only partially explained by growth-rate-correlated proteins. Number of differential expressions (adjusted p-value < 0.01, BH for multiple testing (Benjamini and Hochberg, 1995)) across the KO strain was calculated for each protein, and proteins were grouped into growth-related (green) and non-growth-related proteins (orange) depending on their respective correlation with growth rate (growth-associated: R > 0.2 or R < –0.2, not-growth-associated: –0.2 < R < 0.2). (**D**) Enrichment analysis (hypergeometric test) was performed on the knock-outs that induced aneuploidy using the GO slim gene sets (BP, MF, and CC) (Cherry et al., 2012). Significant terms (adjusted p-value < 0.01) are shown and ranked by significance (decreasing from top to bottom). **(E)** Number of aneuploid knock-outs with and without paralogues. The aneuploid strains with paralogues are grouped into strains where the paralogue is on the aneuploid chromosome (orange) and strains where the paralogue is not on the aneuploid chromosome (green). (**F**) Chromosomes occurring as aneuploid across the KO strains.

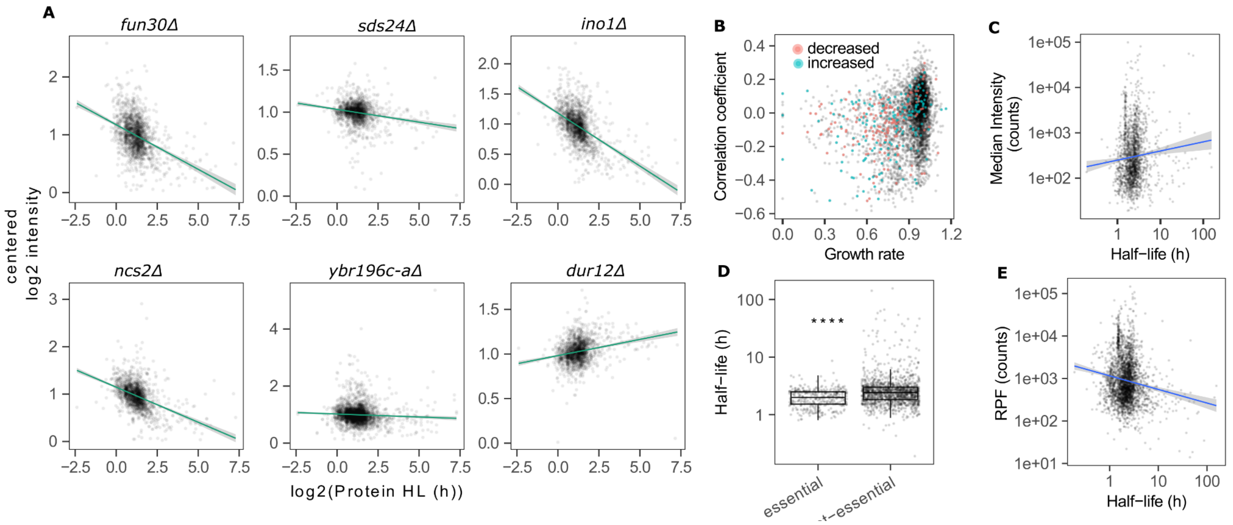


**Figure S4.** (**A**) Half-life-dependent protein-abundance changes for the top 6 features (knock-outs) selected by the elastic net model (*fun30Δ, sds24Δ, ino1Δ, ncs2Δ, ybr196c-aΔ, dur12Δ*). Protein half-lives (Martin-Perez and Villén, 2017) ($log_2$ transformed) are plotted against centred $log_2$ intensites. **(B)** Unspecific half-life-dependent protein-abundance changes are observed across all growth rates and cell sizes. Correlation coefficients (Pearson) were calculated for all strain-wise relationships between protein expression changes and half-lives. Thus, high correlation coefficients indicate a tendency to up-regulate long-lived and down-regulate short-lived proteins. Correlation coefficients are plotted against the growth rate (normalised), and strains with phenotypes characterised by decreased and increased cell size (Cherry et al., 2012) are coloured. **(C)** Relationship between protein abundance (intensities) and half-lives. X-axis is $log_{10}$ transformed. **(D)** Protein half-lives (Martin-Perez and Villén, 2017) are plotted for essential and non-essential genes. Significance (Wilcoxon signed-rank test; **** for p-value ≤ 0.0001) is indicated with asterisks; the first and third quartiles, as well as the median (thick line), are shown with boxplots; whiskers extend to the most extreme data point that is no more than 1.5× the interquartile range from the box. (**E**) Relationship between ribosome-occupancy data (McManus et al., 2014) (counts) and protein half-life data (in h) (Martin-Perez and Villén, 2017). X-axis and y-axis are $log_{10}$ transformed.

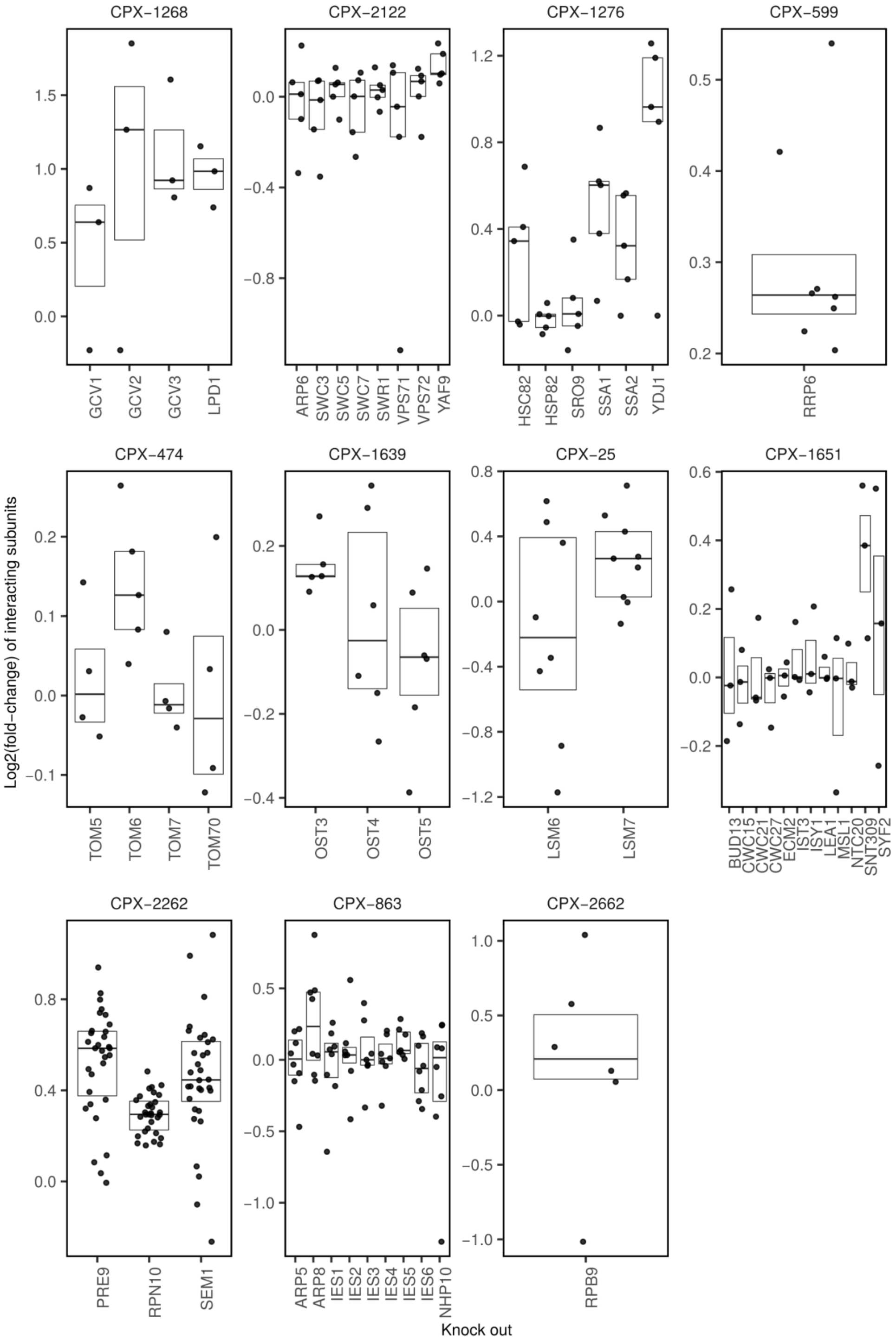


**Figure S5-a. Complexes that result in significant up-regulation upon knock-out of at least one subunit.** Differential expression analysis of the complex subunits was performed against wild-type samples using Wilcoxon signed-rank test and adjusted for multiple correction using BH (Benjamini and Hochberg, 1995). The complexes shown have a significant ($p < 0.05$) difference to the WT samples for at least one knocked-out subunit. Intensities were normalised to the median intensities of all samples (WT and KOs) and $\log_2$ transformed. The first and third quartiles, as well as the median, are shown with boxplots. Complex data were downloaded from the EBI Complex

Portal (Meldal and Orchard, 2018; Meldal et al., 2015, 2019).

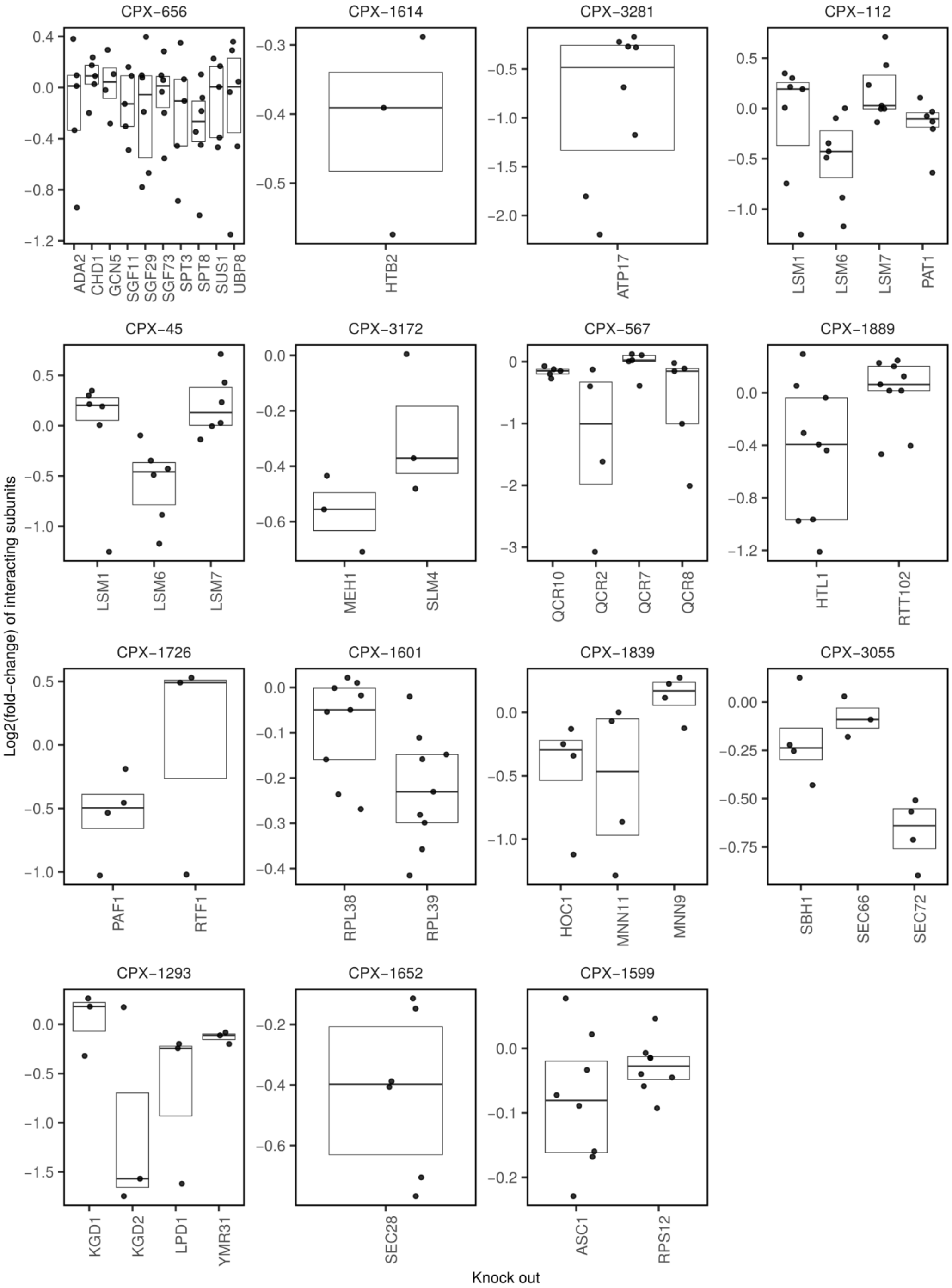


**Figure S5-b. Complexes that result in significant down-regulation upon knock-out of at least one subunit.** Differential expression analysis of the complex subunits was performed against wild-type samples using Wilcoxon signed-rank test and adjusted for multiple correction using BH (Benjamini and Hochberg, 1995). The complexes shown have a significant ($p < 0.05$) difference to the WT samples for at least one knocked-out subunit. Intensities were normalised to the median intensities of all samples (WT and KOs) and $\log_2$ transformed. The first and third quartiles, as well as the median, are shown with boxplots. Complex data were downloaded from the EBI Complex

Portal (Meldal and Orchard, 2018; Meldal et al., 2015, 2019).

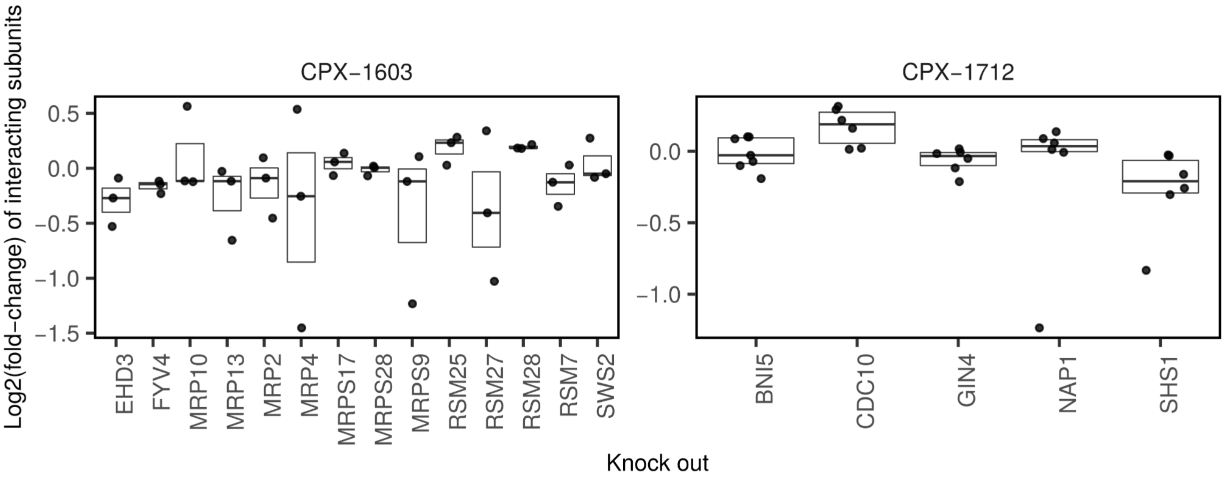


**Figure S5-c. Complexes with subunits that result in up-regulation or down-regulation upon deletion.** Differential expression analysis of the complex subunits was performed against wild-type samples using Wilcoxon signed-rank test and adjusted for multiple correction using BH (Benjamini and Hochberg, 1995). The complexes shown have a significant ($p < 0.05$) difference to the WT samples for at least one knocked-out subunit. Intensities were normalised to the median intensities of all samples (WT and KOs) and $\log_2$ transformed. The first and third quartiles, as well as the median, are shown with boxplots. Complex data were downloaded from the EBI Complex Portal (Meldal and Orchard, 2018; Meldal et al., 2015, 2019).

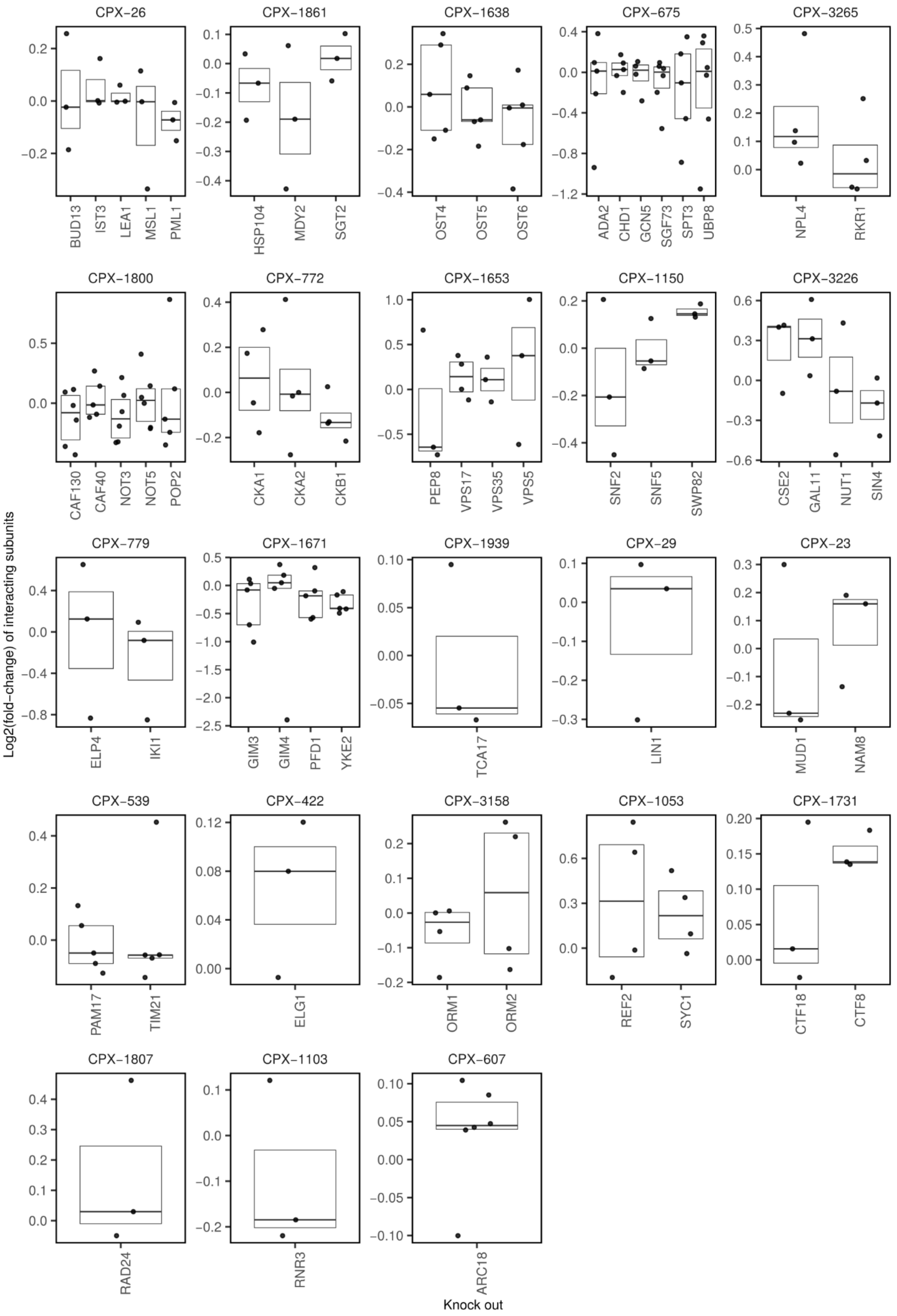


**Figure S5-d. Complexes with subunits that are not affected upon deletion.** Differential expression analysis of the complex subunits was performed against wild-type samples using Wilcoxon signed-rank test and adjusted for multiple correction using BH (Benjamini and Hochberg, 1995). The complexes shown have no significant ($p > 0.05$) difference to the WT samples. The first and third quartiles, as well as the median, are shown with boxplots. Complex data were downloaded from the EBI Complex Portal (Meldal and Orchard, 2018; Meldal et al., 2015, 2019).

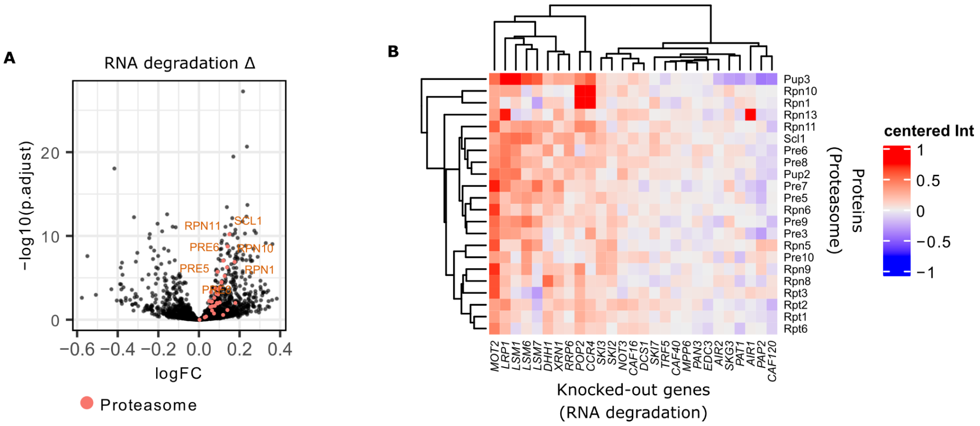


**Figure S6.** (**A) Differential expression for knock-outs involved in RNA degradation.** Knock-out strains were grouped together according to the KEGG RNA degradation term (Kanehisa, 2019; Kanehisa and Goto, 2000) *(CAF120, PAP2, AIR1, PAT1, SKG3, AIR2, EDC3, PAN3, MPP6, CAF40, TRF5, SKI7, DCS1, CAF16, NOT3, SKI2, SKI3, CCR4, POP2, RRP6, XRN1, DHH1, LSM7, LSM6, LSM1, LRP1, MOT2)* and compared to WT samples using the limma package (Ritchie et al., 2015). BH was used for multiple testing (Benjamini and Hochberg, 1995). Proteasome proteins are coloured. $Log_2$ fold changes are shown on the x-axis; adjusted p-values ($-log_{10}$ transformed) on the y-axis. (**B)** RNA-associated knock-outs (Kanehisa, 2019; Kanehisa and Goto, 2000) (horizontally) and significantly changed proteasomal proteins (vertically). Protein intensities were centred and $log_2$ transformed.

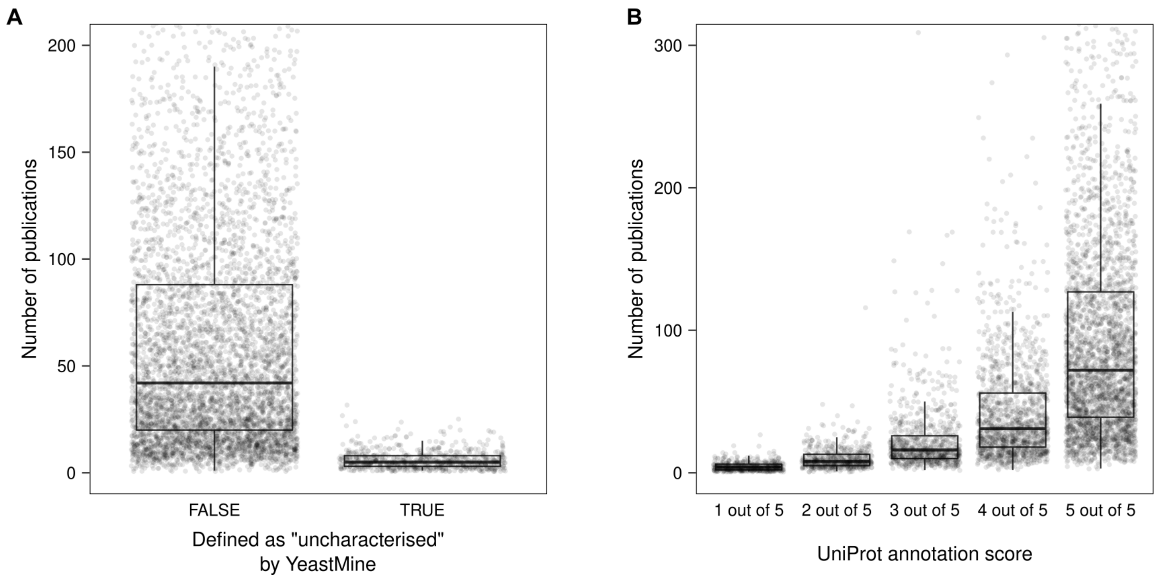


**Figure S7. Many yeast genes are understudied.** (**A**) The number of publications linked to yeast genes according to the Saccharomyces Genome Database (Cherry et al., 2012) is shown with boxplots. YeastMine currently classifies 722 proteins as "uncharacterised". A median of 5 publications can be mapped to these, compared to a median of 42 publications for the remaining genes. (**B**) Same data as in (A), but genes were divided based on the annotation score assigned to each gene by UniProt. The 2,913 best-annotated yeast genes (5 out of 5) have a median of 103 publications each, whereas the 468 worst-characterised genes (1 out of 5) have a median of 4 publications.

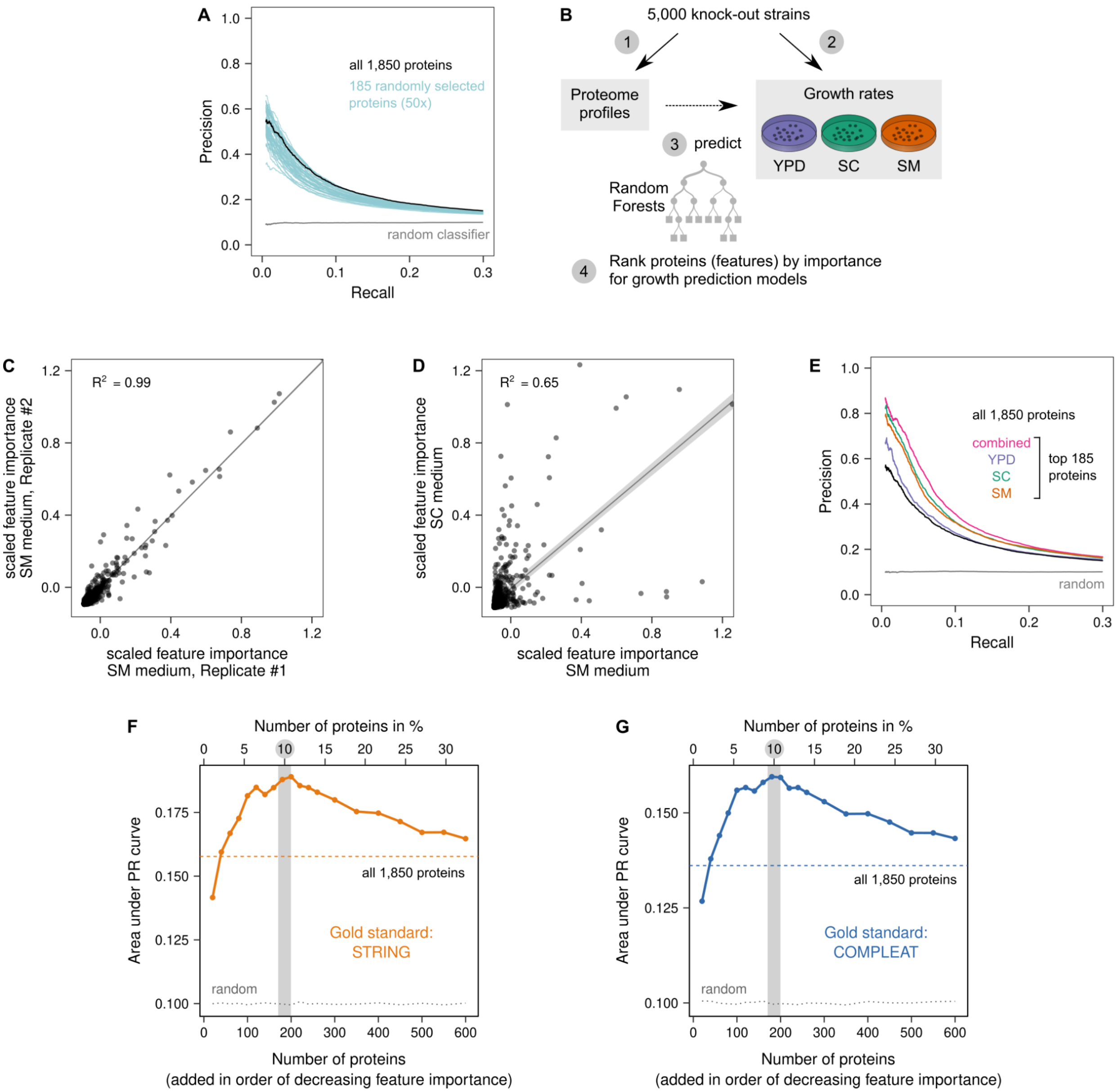


**Figure S8. Feature selection for proteome profile similarity assessment.** (**A**) Similarities between knock-out strains are challenging to define in a high-dimensional space consisting of 1,850 proteins. We subjected 50 randomly selected groups of 185 proteins (10%) to precision–recall (PR) analyses using the STRING gold standard. Although this is only a miniscule fraction of the theoretically possible $5 \times 10^{259}$ 185-protein combinations, several of these randomly selected subsets identify functionally related knock-out genes with higher precision than a PR analysis using all proteins. This indicates that the high dimensionality of these data is a problem ("curse of dimensionality") and that functional predictions could be improved by selecting an optimal subset of proteins (feature selection). (**B**) A common strategy for feature selection in data science is the use of random forests (RFs), which offer a straightforward way to assess the importance of each feature for a regression model. We measured the growth rates of the KO strains in three growth media (YPD, SM, SC). We then trained RF regression models to predict these KO strain growth rates from the abundances of the 1,850 quantified proteins. The importance of each feature (protein) for these predictions was extracted from the RF models. (**C**) High technical reproducibility of determining feature (i.e. protein) importance in this manner is shown using two replicate RF models. Feature importances extracted from two replicate analyses are highly similar. (**D**) In contrast, feature importances differ considerably between RF models predicting growth in different growth media. As an example, this plot compares feature importances for predicting growth in SM and SC media. (**E**) PR analysis similar to (A), but using the 185 proteins with the highest feature importances for growth-rate predictions. For all three predictions this subset outperforms the use of all 1,850 proteins. Notably, performance can be improved further by combining feature importances across the three growth media, which is achieved by ranking proteins based on the minimal scaled importance they achieved in any RF model. (**F**) To determine the optimal number of features (proteins) to select in this way, proteins were ranked by feature importance (across all three growth media) and a series of PR analyses was performed. The plot shows the areas under the PR curves. We find that performance increases as more proteins are added, plateaus around 185 proteins (10% of the 1,850 available proteins), and then decreases again. This suggests that to compare proteome profile similarities of KO strains it is best to consider only the 10% of proteins with the highest feature importance. (**G**) Same as (F) but using a different gold standard (COMPLEAT) as a reference for the PR analyses, with a very similar outcome.

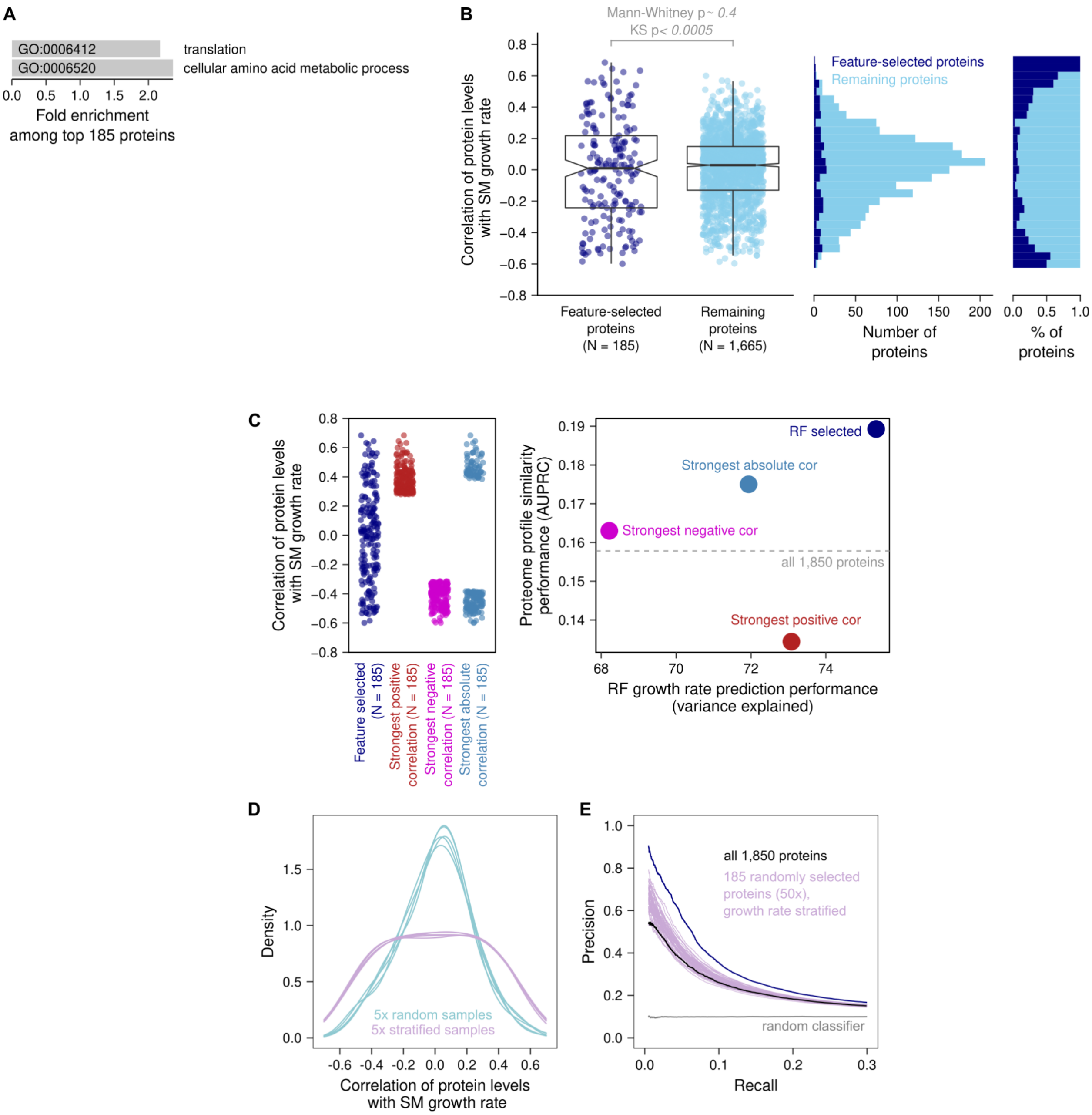


**Figure S9. Selected features are stratified by growth-rate dependence.** (**A**) The 185 proteins selected based on their importance for growth-rate prediction by the Random Forest (RF) are enriched in two GO slim *biological processes* (FDR 0.05), relative to the all 1,850 detected proteins. This was detected by a statistical overrepresentation test using Panther (Mi et al., 2016). (**B**) To determine if the RF simply enriches for proteins whose expression is determined by the growth rate, we analysed the correlation of protein fold changes with the growth rates of the KO strains. Interestingly, we find that the 185 selected proteins are on average not correlated more strongly with the growth rate than the remaining 1,665 proteins (Mann–Whitney p-value ~ 0.4), but the distribution of these correlations is significantly different (Kolmogorov–Smirnov p-value < 0.0005). This can also be seen in the histogram: the selected proteins are uniformly distributed across the correlation range. This means that the RF does enrich proteins that are either correlated or anti-correlated with the growth rate, but not exclusively. In effect, the RF-based feature selection therefore results in a protein selection that is stratified by growth-rate dependence. (**C**) To better understand the relationship between growth-rate dependence and feature importance, we compared the 185 feature-selected proteins to the performance of 185 proteins that most strongly correlate with growth rates, either positively, negatively, or on an absolute scale. It shows that the feature-selected proteins are the best subset for both growth-rate prediction and assessment of proteome profile similarities of KOs. Intriguingly, proteins that positively correlate with the growth rate work well to predict growth rates, but are a poor choice for assessing proteome profile similarity. Proteins that negatively correlate with growth rates can connect functionally related KOs relatively well, but are poorer features for growth-rate prediction. (**D**) We use a stratified sampling strategy, where we select 185 proteins randomly but in a way that they evenly cover the whole range of growth-rate dependence, from strong positive to strong negative. The density plot shows five such samples in comparison to five completely random 185-protein samples. (**E**) We then assess if this "stratified sampling approach" could be a suitable approach for feature selection for protein function prediction. Out of 50 random stratified samples, 43 outperform a precision–recall (PR) analysis based on all proteins. This compares favourably to the 5 of 50 for totally random samples (Figure S8).

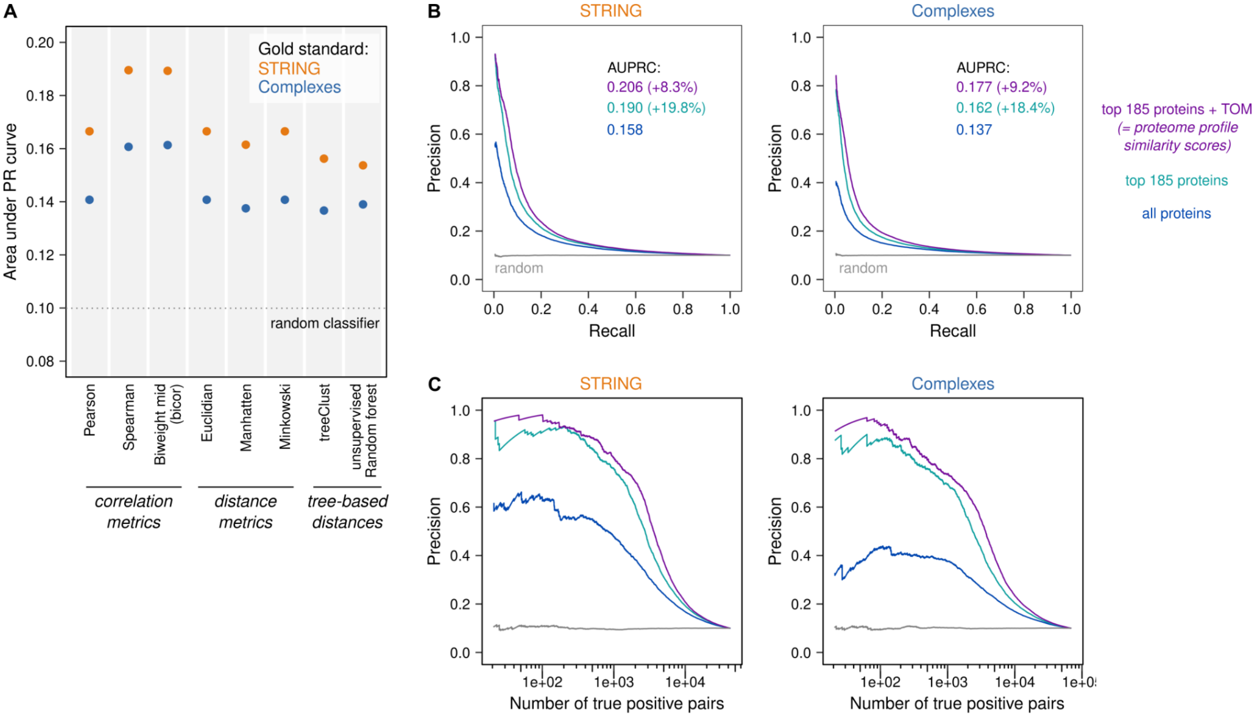


**Figure S10. Calculation of proteome profile covariation between KO strains.** (**A**) We compared a range of correlation and distance metrics for how well they identify profile covariation across the ~5k yeast KO strains, on the basis of the 185 pre-selected proteins. Performance was assessed with precision–recall (PR) analyses using two different gold standards (STRING and protein complexes from Compleat (Vinayagam et al., 2013)). The areas under the PR curves are shown. Optimal performance is observed for two types of robust correlation metrics, Spearman's correlation and biweight midcorrelation, with the latter becoming our preferred choice as it can be calculated more efficiently (19 min vs 10 sec using R on a standard PC). (**B**) A topological overlap measure (TOM) further improves the precision with which KO strains of functionally related genes can be linked. The PR analyses use either STRING (left panel) or COMPLEAT (right panel) gold standards, insets show the area under the PR curves (AUPRC). The PR curves show how well the Y5k KO strains can be linked by biweight midcorrelation. Feature selection improves performance by 18.4–19.2% compared to correlating all 1,850 quantified proteins. Taking into account the topology of the resulting correlation network helps to remove false-positive links and thereby improves performance by an additional 8.3–9.2%. These TOM-modified, biweight midcorrelations of the 185 selected proteins constitute our profile covariation scores. (**C**) Same PR curves as in (B), but expressing the recall as absolute number (rather than fraction) of true positive pairs. This shows that the performance improvement brought by feature selection and topological overlap measure is particularly pronounced for the first few thousand gene–gene associations.

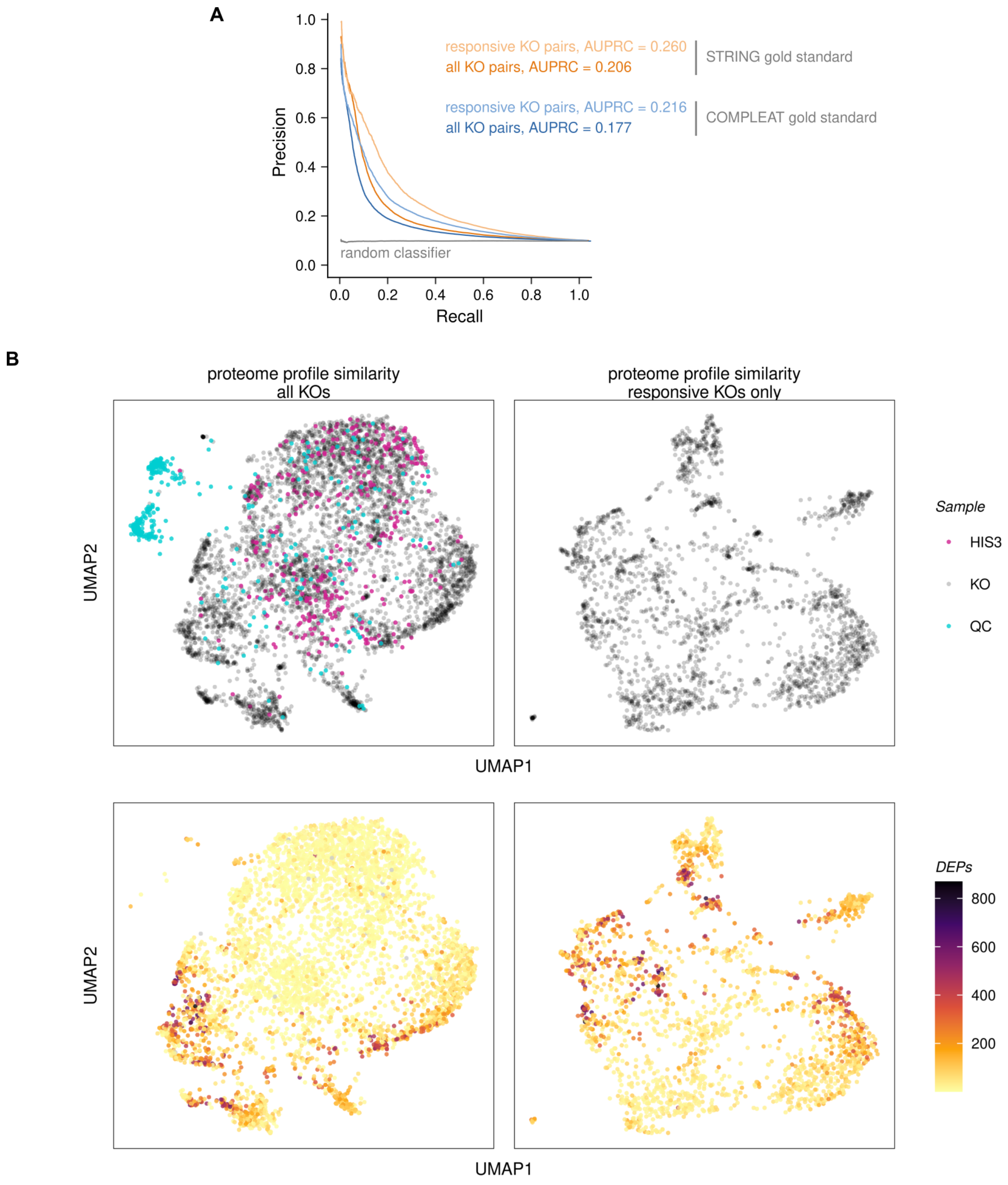


**Figure S11. Responsive KOs are better suited for function prediction by proteome profile covariation.** (**A**) Precision–recall (PR) curve showing that focussing the analysis on the 2,290 "responsive" KO strains strongly improves performance (STRING AUPRC 0.26 vs 0.20 for all KOs, an improvement of ~26%). This means the proteome profiles of responsive KOs can be compared more accurately and will therefore lead to better gene-function predictions. A "responsive" strain is defined here as having more differentially expressed proteins (DEPs) than the median strain. (**B**) UMAP plots of proteome profile similarities. Each point is a KO strain and the proximity between points indicates how similar their proteome profiles are. Measurements for all 5,463 KO strains, wild-type, and QC controls are shown on the left panel; the 2,290 "responsive" KO strains only are shown on the right. Top: QC controls group together and most wild-type (HIS3) strains are located at the top of the map. Bottom: The number of DEPs per KO is highlighted. KOs with many or few DEPs, respectively, tend to group together. The top area containing the wild-type strains is generally devoid of DEPs. Even when those KOs are discarded from the analysis (right panel, responsive KOs only) there are areas with higher and lower DEP levels.

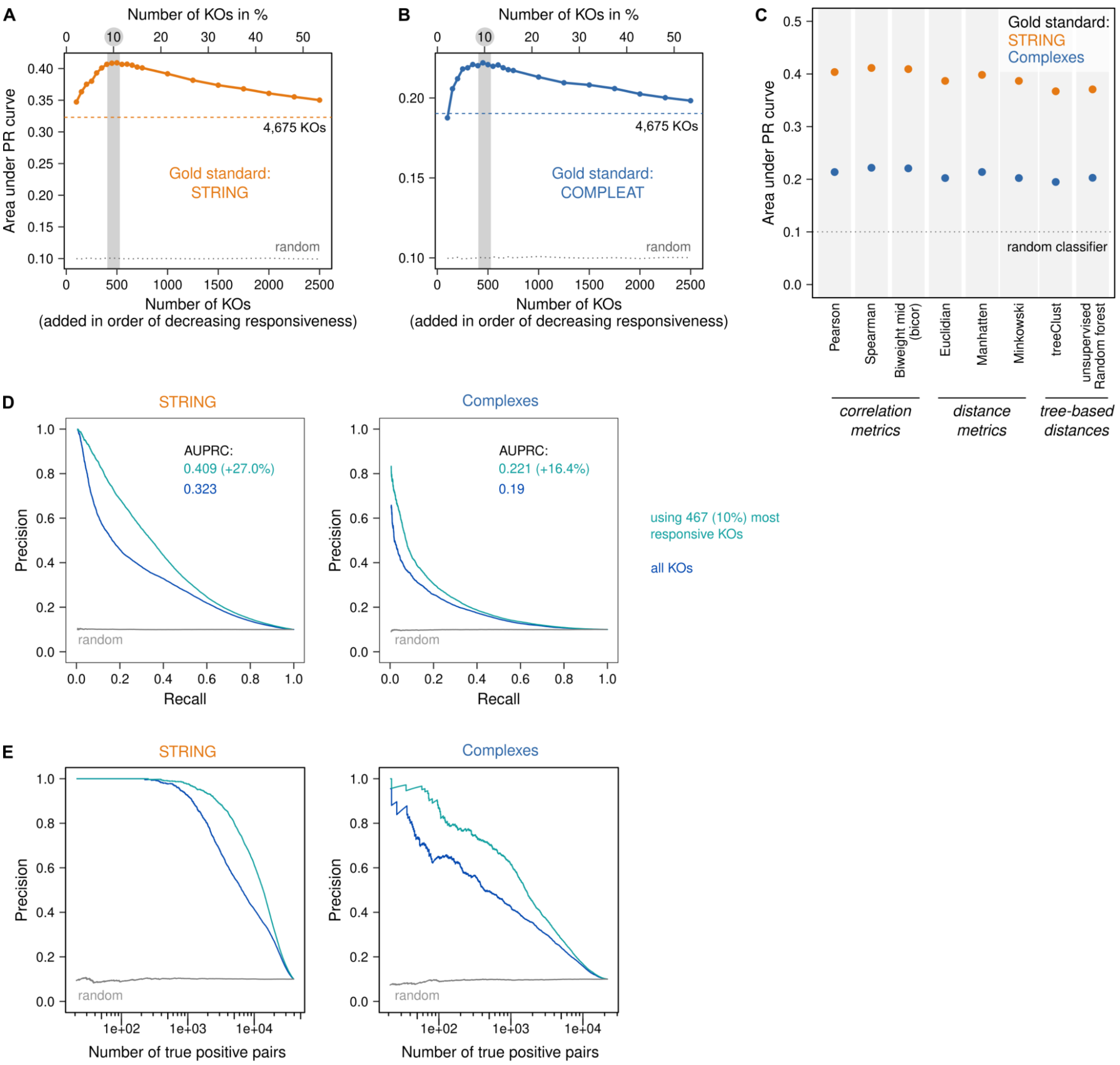


**Figure S12. Determination of protein covariation scores.** (**A**) Feature selection for the protein covariation analysis. We reasoned that "responsive" KO strains may be best suited to identify co-regulated proteins. To test this, proteins were ranked by "responsiveness", which was defined as the number of differentially expressed proteins identified by limma analysis. We then performed a series of precision–recall (PR) analyses, starting with the 100 most responsive strains and gradually including more strains up until using all 4,675 KO strains that had been included in the limma analysis. The plot shows the areas under the PR curves. We find that performance increases initially as more KOs are added, but plateaus around 450–500 strains and then decreases again. We therefore decided to use the top 467 (10%) most responsive KO strains to measure protein co-regulation. (**B**) Same as (A) but using a different gold standard (protein complexes from Compleat (Vinayagam et al., 2013)) as a reference for the PR analyses, with a very similar outcome. (**C**) We compared a range of correlation and distance metrics for how well they identify co-regulated proteins across the 467 pre-selected KO strains. Performance was assessed in PR analyses using two different gold standards (STRING and Compleat complexes) as a reference. The areas under the PR curves are shown. Optimal performance is observed for two types of robust correlation metrics, Spearman's correlation and biweight midcorrelation, with the latter becoming our preferred choice as it can be calculated more efficiently. (**D**) The feature (KO) selection process improves the precision with which co-regulated proteins can be identified. The PR analyses use either STRING (left panel) or Compleat (right panel) gold standards, insets show the area under the PR curves (AUPRC). The analysis used biweight midcorrelation. Feature selection improves performance by 16.4–27% compared to correlating all ~5k yeast strains. Note that in contrast to the proteome-profile-covariation network of KOs, taking into account the topology of the protein covariation network did not improve performance further and was therefore omitted. Consequently, the biweight midcorrelations across the 476 selected KO strains constitute our protein covariation scores. (**E**) Same PR curves as in (D) but expressing the recall as absolute number (rather than fraction) of true positive pairs.

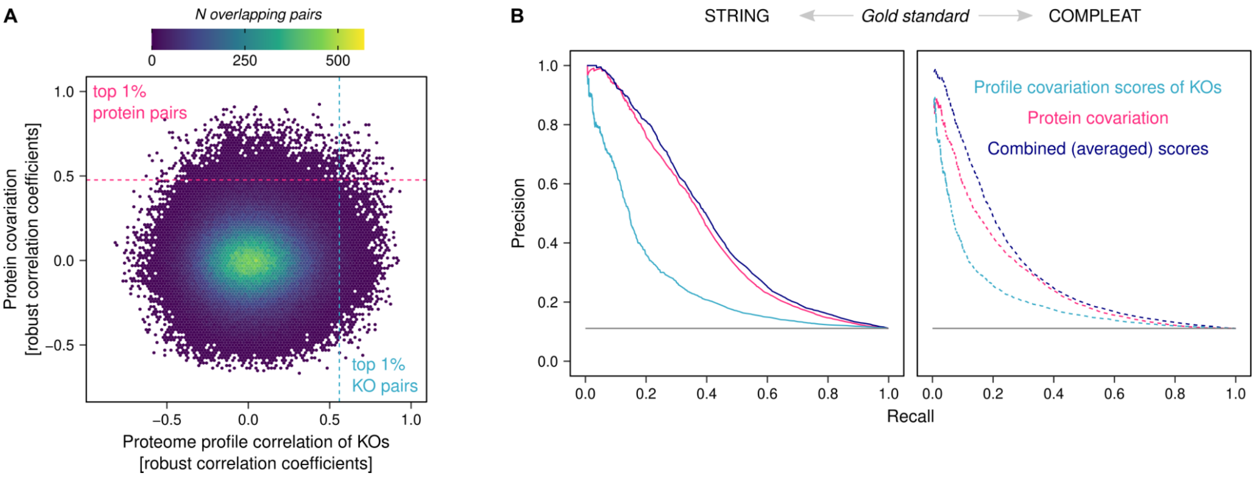


**Figure S13. Proteome profile covariation of KOs and protein covariation across KOs capture complementary gene–gene associations.** (**A**) Robust correlation coefficients of pairwise associations that were detected either by proteome profile similarity of KO strains or by protein covariation are plotted against each other. There is no common trend and the top 1% associated pairs by each approach overlap only marginally. (**B**) We tested if the two approaches are not just complementary quantitatively (i.e. capturing more genes and associations together than each would on its own), but also qualitatively. Indeed, when profile and protein covariation scores are scaled and averaged for the pairs that they both cover, the combined scores outperform individual ones in a precision–recall analysis.

**Tabe S1:** Lower and upper m/z limits of precursor isolation windows in the applied variable window SWATH-MS method.

| Lower m/z | Upper m/z |
|---|---|
| 399.5 | 410.4 |
| 409.4 | 420.3 |
| 419.3 | 429.7 |
| 428.7 | 439.6 |
| 438.6 | 448.9 |
| 447.9 | 458.8 |
| 457.8 | 468.7 |
| 467.7 | 478.1 |
| 477.1 | 488.5 |
| 487.5 | 499 |
| 498 | 509.4 |
| 508.4 | 519.9 |
| 518.9 | 530.9 |
| 529.9 | 541.9 |
| 540.9 | 552.8 |
| 551.8 | 564.4 |
| 563.4 | 575.9 |
| 574.9 | 588 |
| 587 | 600.1 |
| 599.1 | 612.2 |
| 611.2 | 625.4 |
| 624.4 | 639.2 |
| 638.2 | 652.9 |

| | |
|---|---|
| 651.9 | 667.2 |
| 666.2 | 681.5 |
| 680.5 | 696.9 |
| 695.9 | 712.3 |
| 711.3 | 728.8 |
| 727.8 | 747.5 |
| 746.5 | 766.2 |
| 765.2 | 786 |
| 785 | 806.9 |
| 805.9 | 828.9 |
| 827.9 | 854.8 |
| 853.8 | 882.8 |
| 881.8 | 915.3 |
| 914.3 | 957.6 |
| 956.6 | 1015.4 |
| 1014.4 | 1098.4 |
| 1097.4 | 1249.7 |

**Table S2.** Timing and mobile-phase composition for the applied non-linear chromatography gradient.

| Time (min) | Mobile phase (% A) |
|---|---|
| initial | 97 |
| 0.86 | 92.9 |
| 2.42 | 88.8 |
| 5.53 | 84.7 |
| 9.38 | 80.6 |
| 13.02 | 76.4 |
| 15.48 | 72.3 |
| 17.27 | 68.2 |
| 19 | 60 |
| 20 | 20 |
| 20.5 | 20 |
| 21.5 | 97 |
| 27.5 | 97 |

**Table S3:** Knock-out strains selected for the prediction of ribosomal occupancy (McManus et al., 2014). The top 15 strains selected by the elastic net model are listed below with corresponding feature importance. Feature selection was done with the varimp function within the caret package (Kuhn, 2008).

| Gene name | Feature importance |
|---|---|
| *elp4Δ* | 100 |
| *spt10Δ* | 89.19691 |
| *mot2Δ* | 88.79057 |
| *ubp3Δ* | 86.14195 |
| *def1Δ* | 84.50996 |
| *yjl175wΔ* | 80.81533 |
| *fyv4Δ* | 80.60749 |
| *npr3Δ* | 78.62733 |
| *ykl169cΔ* | 74.90061 |
| *ino1Δ* | 71.6569 |
| *get1Δ* | 69.56296 |
| *blm10Δ* | 66.9569 |
| *bar1Δ* | 64.10587 |
| *seh1Δ* | 59.31983 |
| *ecm30Δ* | 58.42478 |

**Table S4**: Knock-out strains selected for the prediction of protein half-life (Martin-Perez and Villén, 2017). The top 15 strains selected by the elastic net model are listed below with corresponding feature importance. Feature selection was done with the varimp function within the caret package (Kuhn, 2008).

| Strain | Feature importance |
|---|---|
| *fun30Δ* | 100 |
| *sds24Δ* | 96.3988 |
| *ino1Δ* | 95.01005 |
| *ncs2Δ* | 88.48427 |
| *ybr196c-aΔ* | 84.96767 |
| *dur12Δ* | 82.29651 |
| *gim4Δ* | 72.66398 |
| *mot2Δ* | 70.20363 |
| *ybr220cΔ* | 69.28264 |
| *vps51Δ* | 59.81237 |
| *get1Δ* | 59.42979 |
| *rps11bΔ* | 56.14677 |
| *bud25Δ* | 55.60507 |
| *ydr048cΔ* | 53.71978 |
| *sbe2Δ* | 53.65612 |

**Table S5**: Proteins that are up-regulated upon deletion of their paralogue. The annotations of paralogues were downloaded from the Yeast Gene Order Browser (Byrne and Wolfe, 2005).

| Knock-out strain | Up-regulated protein | Fold change | Adjusted p-value |
|---|---|---|---|
| *rps14aΔ* | Rps14b | 4.085177 | 6.7E-34 |
| *rpl4aΔ* | Rpl4b | 2.72258 | 4.6E-04 |
| *rpl8aΔ* | Rpl8b | 1.310114 | 6.1E-04 |
| *rps29bΔ* | Rps29a | 1.372421 | 2.3E-05 |
| *rps1aΔ* | Rps1b | 1.534089 | 1.2E-14 |
| *rpl6bΔ* | Rpl6a | 1.36172 | 2.4E-04 |
| *rnr4Δ* | Rnr2 | 6.540719 | 5.7E-119 |
| *rpl7aΔ* | Rpl7b | 4.987068 | 2.3E-48 |
| *ura7Δ* | Ura8 | 1.691639 | 1.6E-09 |
| *rps9bΔ* | Rps9a | 5.445963 | 4.4E-03 |
| *cit1Δ* | Cit2 | 2.337526 | 5.5E-03 |
| *rps7aΔ* | Rps7b | 1.858272 | 6.7E-13 |
| *rpl16bΔ* | Rpl16a | 2.150493 | 5.7E-46 |
| *rpl14aΔ* | Rpl14b | 1.772908 | 2.6E-06 |
| *rpl8bΔ* | Rpl8a | 2.291356 | 2.7E-13 |
| *tif4631Δ* | Tif4632 | 1.952875 | 4.1E-03 |
| *rps29aΔ* | Rps29b | 1.473908 | 7.4E-06 |
| *rpl6aΔ* | Rpl6b | 2.249223 | 8.2E-39 |
| *sse1Δ* | Sse2 | 2.356487 | 8.0E-06 |
